# Inferring Spatially Varying Bending Stiffness of Biopolymers with Deep-Learning Approach

**DOI:** 10.1101/2025.06.12.659266

**Authors:** Changbeom Hong, Chan Lim, Jaeho Oh, Jong-Bong Lee, Euddeum Eojin Jeong, Joo-Yeon Yoo, Won Kyu Kim, Jae-Hyung Jeon

**Affiliations:** Department of Physics &, Pohang University of Science and Technology, Pohang 37673, Republic of Korea; Institute for Theoretical Science, Pohang University of Science and Technology, Pohang 37673, Republic of Korea; Department of Physics, Pohang University of Science and Technology, Pohang 37673, Republic of Korea; Department of Life Sciences, Pohang University of Science and Technology, Pohang 37673, Republic of Korea; Korea Institute for Advanced Study, Seoul 02455, Republic of Korea; Department of Physics and Astronomy, Seoul National University, Seoul 08826, Republic of Korea; Asia Pacific Center for Theoretical Physics, Pohang 37673, Republic of Korea

**Keywords:** biopolymers, persistence length, spatially varying bending stiffness, deep-learning, transformer model

## Abstract

The bendability of biopolymers, including DNA, actin filaments, and microtubules, is crucial to diverse biological processes, such as cell motility and cytoskeletal organization. While conventional polymer models like the worm-like chain model assume uniform bending stiffness along the contour, biopolymers in reality often attain spatially varying bending stiffness as a result of complex cellular interactions. For instance, the bending stiffness of actin filaments and microtubules varies in response to associated binding proteins or chemical modifications. Despite its biological implications, measuring the position-dependent persistence length along polymer contours remains challenging. Here, we present a deep-learning-based method that quantitatively predicts spatially varying bending stiffness of biopolymers. Our framework segments a polymer chain into short overlapping fragments, predicts local persistence lengths using a deep-learning model, and reconstructs full profiles. Using simulated data, we demonstrate that our approach achieves high accuracy and robustness even under data-scarce conditions. Applied to various biological systems, our method reveals a spatially varying stiffness from tip to base in filopodia as well as spatially heterogeneous stiffness in microtubules. Additionally, we investigate the internal mechanism of our deep-learning model and show that the model utilizes multiple physics-related features for persistence length estimation while adaptively adjusting its attention based on polymer stiffness.

## I. INTRODUCTION

Biopolymers such as DNA, actin filaments, and microtubules play crucial roles in a wide range of biological processes, including gene regulation, molecular transport, cell motility, and cytoskeletal formation [1– 3]. Accordingly, their mechanical and structural properties have been extensively studied. Traditional theoretical frameworks, such as the worm-like chain (WLC) model, typically assume that biopolymers are mechanically uniform along the polymer contour. However, it has been revealed that many biopolymers often exhibit spatial heterogeneities in their stiffness, driven by interactions with complex cellular environments. For example, in double-stranded DNA, local denaturation, frequently triggered by thermal fluctuation or molecular interactions with single-stranded binding proteins, significantly reduces persistence length, and it is closely related to the DNA sequence [4–6]. Moreover, damaged DNA sites become more flexible, enabling repair proteins to recognize and bind these regions based on mechanical cues [7]. Actin filaments show similar heterogeneity; ADF/cofilin proteins reduce the persistence length of the actin filament, which can lead to enhanced filament fragmentation [8–10]. Likewise, microtubules exhibit variable bending stiffness in response to specific proteins or drug molecules, as well as due to chemical modifications such as acetylation and deacetylation [11, 12].

Despite its biological significance, quantifying the spatially varying persistence length along a polymer contour remains a challenging task. Although several studies have developed methods to extract mechanical properties using the force-response relation or the fluctuation of the polymer, these methods are inapplicable to probe local mechanical properties of biopolymers [13–16]. In contrast, Wong *et al*. developed a microrheology technique to probe local mechanical properties of biopolymers, while Bangalore *et al*. measured the flexibility of damaged DNA sites by analyzing turning angles at the lesion sites [7, 17]. However, these approaches primarily investigate local changes in stiffness and do not provide a comprehensive profile of mechanical properties along the full polymer contour. Valdman *et al*. developed a method to quantify heterogeneous elasticity using a modified WLC model combined with a sliding window method, but this approach is applicable only to relatively stiff polymers [18].

In parallel, deep-learning techniques have rapidly advanced, yielding notable contributions across a wide range of disciplines. In biophysics, deep-learning has played a pivotal role in resolving long-standing challenges. A prominent example is AlphaFold, which achieved a breakthrough in protein structure prediction [19–21]. Beyond protein folding, deep-learning approaches have also facilitated a deeper understanding of complex biological systems. Notably, the AnDi Challenge demonstrated the effectiveness of these methods in elucidating intracellular transport dynamics by enabling the analysis of diverse anomalous diffusion models from single-particle tracking data [22]. In the field of polymer physics, deep-learning methods have also been employed to address a range of problems, including knot type classification, phase & phase transition prediction, and estimation of polymer properties such as electron affinity and ionization potential [23–25]. In line with these efforts, we propose a deep-learning framework that infers the persistence length of biopolymers from two-dimensional chain conformation data, which can be obtainable from microscopy images.

In this study, we develop a deep-learning framework for accurately predicting the profile of spatially varying persistence lengths of biopolymers. Our approach employs a transformer-based deep-learning architecture to estimate local persistence lengths. This allows the model to learn mutual dependencies between monomers, enabling accurate prediction of local persistence lengths, as well as reconstruction of the full persistence length profile along the polymer contour. Using simulated polymer backbone configurations, we benchmark our method against conventional methods based on statistical estimators and demonstrate superior performance in estimation accuracy. Our approach exhibits strong robustness to limited data in comparison with conventional methods, making it particularly suitable for real-world experimental conditions. We further apply our framework to a variety of biological systems, including DNA, actin filaments, in vivo filopodia, and intracellular microtubules, uncovering meaningful biological insights such as a tipto-base stiffness gradient in filopodia and local stiffness heterogeneity in microtubules. Finally, we investigate the inner workings of our deep-learning model to provide physics-informed insights into its operation. This analysis reveals that the model leverages multiple physically relevant quantities for persistence length estimation and adjusts attention strategies based on polymer stiffness.

Our study is organized as follows. Section II constructs the theoretical foundation to describe spatially varying bending stiffness based on the WLC model and a deeplearning framework for quantifying the heterogeneous persistence length profiles. In section III, we benchmark the performance of our proposed method against conventional approaches on simulated polymer conformations characterized with both uniform and spatially varying persistence lengths. Section IV demonstrates the applicability of our approach to various biological systems, including DNA, actin filaments, filopodia, and microtubules, and offers their biophysical interpretations. Finally, in section V, we investigate the latent representations and self-attention mechanisms of our deep-learning model, providing physically interpretable insights into its inner workings.

## II. DEVELOPMENT OF DEEP-LEARNING METHOD

In this section, we explain a mathematical framework for describing semiflexible polymers with a spatially varying bending stiffness and outline our deep-learning method for prediction of the persistence length profile from the polymer’s two-dimensional coordinate data.

### A. Worm-like chain model

Polymers are long chains consisting of repeating molecular units. Due to their connectivity and thermal fluctuations, they exhibit a distinct, rich statistical behavior, which is quite different from monomeric systems, depending on their stiffness, contour length, and environmental conditions. A central challenge in polymer physics is to quantitatively describe the morphological properties of polymers, which is both mathematically tractable and physically meaningful. The worm-like chain model (also known as the Kratky–Porod model in its discrete form) serves as a foundational framework for describing the semiflexible nature of many biopolymers, including DNA, actin filaments, and microtubules [13, 14, 26, 27]. It models the polymer as a continuous and differentiable space curve **r**(*s*) with an energy cost associated with bending (curvature change), parameterized by the arc length *s* ∈ [0, *L*], where *L* is the contour length. The local orientation is given by the unit tangent vector **t**(*s*) = *d***r**(*s*)*/ds*, with |**t**(*s*)| = 1. The bending energy is

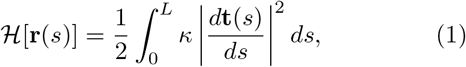

where *κ* is the bending modulus. Thermal fluctuations lead to configurational statistics governed by the Boltzmann distribution ∈ exp[−ℋ */*(*kBT*)]. From Eq. (1), the Boltzmann-weighted ensemble average of tangent vectors yields the well-known exponential correlation function [28]

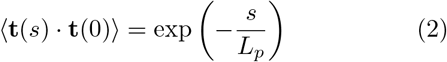

Where

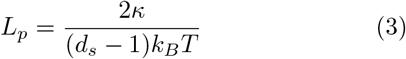

is the persistence length, proportional to the bending modulus *κ*, indicating the characteristic lengthscale of a polymer chain’s directional memory. The *ds* refers to the spatial dimension of the embedding space. A related important physical observable is the mean-squared endto-end distance,

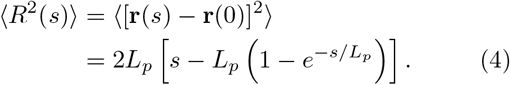

These two relations, Eqs. (2) and (4), are often employed to extract the value of the persistence length *Lp* from an ensemble of polymer chain conformations.

More generally, the bending modulus *κ* can be position dependent, and so is *Lp*. Therefore, the bending energy is

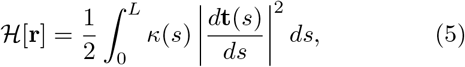

which yields

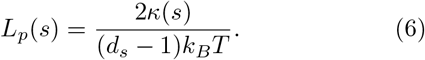

This characterizes the position-dependent stiffness profile of a polymer (i.e., the profile of a persistence length along the contour).

In this study, we focus on the two-dimensional discrete WLC model (i.e., *ds* = 2), reflecting the dimensionality of conventional biopolymer imaging techniques, which yield 2D projections [29, 30]. Moreover, many trajectory-tracking algorithms process discretized 2D coordinates extracted from such images [31, 32]. Accordingly, we estimate *Lp* in 2D and convert it to the corresponding 3D value by dividing by a factor of 2.

All lengths are expressed in units of *a* throughout the study, in which we rescale experimental data with a polymer segment length *a*. Thus, the continuous arc-length parameter, *s* ∈ [0, *L*], is replaced by an integer index *n* ∈ [0, *Nm*] with *Nm* = *L/a*. Since we set *a* = 1, *L* denotes both the total contour length and the number of bonds in the discrete chain, which consists of *L* + 1 monomers at positions 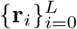 (see Appendix A for the discrete WLC formulation).

### B. Deep-learning method

In Fig. 1, we present our prediction framework based on a deep-learning (DL) model to quantify the profile of local persistence lengths of polymers. The core of our approach is a deep neural network designed to extract spatially resolved elastic properties directly from polymer chain conformation. Our framework consists of three steps: segmentation, pre-estimation, and reconstruction.

**FIG. 1.**
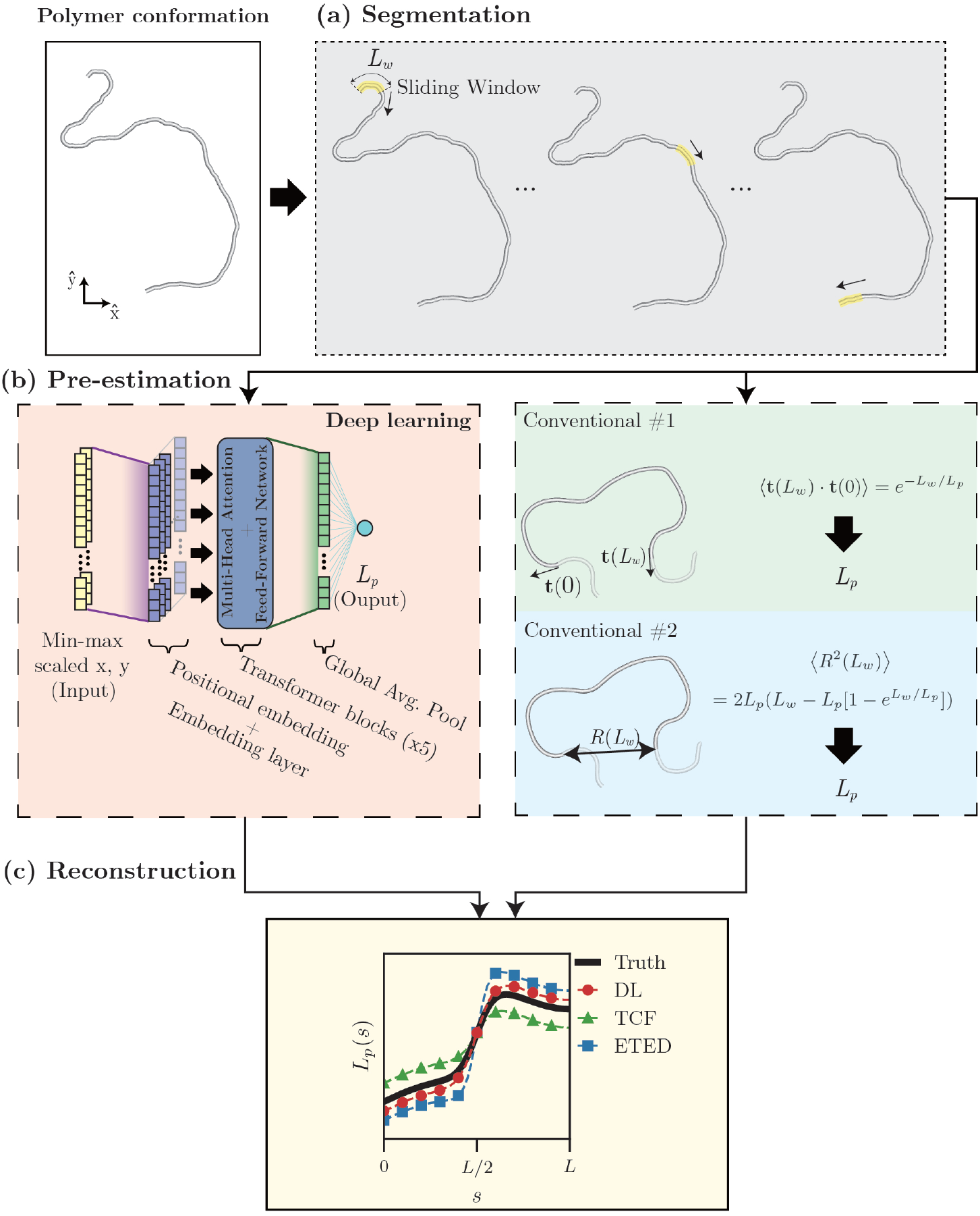
Workflow of our prediction model to quantify the polymer’s position-dependent persistence length profile, *L*_*p*_(*s*), as a function of arc length *s*. (a) Segmentation: a polymer chain is divided into overlapping segments using a sliding window of length *L*_*w*_ along the contour. (b) Pre-estimation: *L*_*p*_ of each segment is estimated by either a deep-learning model or conventional statistical methods, respectively. (c) Reconstruction: the full persistence length profile *L*_*p*_(*s*) is reconstructed and compared with those from the conventional methods.

1. The segmentation step: As illustrated in Fig. 1(a), a polymer chain of length *L* is divided into *L* − *Lw* + 1 overlapping segments of length *Lw* = 24 (*Lw < L*). Therefore, we use the sliding window of length *Lw*, which moves forward one monomer (*a*) subsequently.
2. The pre-estimation step: A pre-trained DL model estimates *Lp* for each overlapping segment [Fig. 1(b)]. The detailed procedure is provided in Appendix B. For comparison, the conventional statistical estimators, the tangent-tangent correlation (Eq. (2)) and the end-to-end distance (Eq. (4)), are computed separately.
3. The reconstruction step: As depicted in Fig. 1(c), the profile of local persistence lengths is reconstructed in terms of mean *Lp* per window as a function of the contour *s* via the sliding window averaging (see Appendix B for further information). For comparison, the results from the tangent-tangent correlation and the end-to-end distance are also presented.

#### 1. Design of model architecture

Several studies have shown that the coordinate representation of polymers, along with appropriate scaling, provides valuable features for predicting polymer properties compared to other geometric quantities such as pairwise displacements, bond angles, and dihedral angles [23, 24, 33, 34]. Motivated by these studies, our DL model takes input data in the form of min-max scaled *x* and *y* coordinates of a polymer segment and outputs its estimated *Lp*, as shown in Fig. 1(b). Many existing algorithms for tracing biopolymers from experimental images yield two-dimensional coordinate data, making our model readily applicable to experimental data [31, 32]. In addition, we apply an additional preprocessing step prior to input: each polymer segment is initially aligned along the positive *x*-axis before scaling. As bending of a polymer is accumulated along the contour, the initial orientations of polymer segments can differ significantly. As a result, this alignment step enables the chain coordinate data to be consistently fed into the model. Subsequently, the *x* and *y* coordinates of the rotated polymer segment are independently scaled by a min-max scaler with a feature range between −1 and 1. The detailed procedure and formulation for these preprocessing steps are provided in Appendix C 1.

To design an effective model architecture, we have examined the performance of various architectures, including feed-forward neural networks (FFNNs), convolutional neural networks (CNNs), recurrent neural networks (RNNs), and transformers, and found that the transformer-based architecture outperforms the others. Given these preliminary investigations, we design our deep-learning model based on the encoder component of the transformer architecture proposed in Ref. [35]. The transformer is designed to capture short- and long-range dependencies between input components and has demonstrated state-of-the-art performance in a wide range of applications [20, 21, 36, 37]. In the context of polymers, short-range dependencies are related to local bending between consecutive monomers, while long-range dependencies are related to the length scale over which the polymers maintain their directional coherence, as quantified by the persistence length. These dependencies suggest that a transformer-based architecture is well-suited for our purpose. Furthermore, self-attention scores provide direct insight into how the attention mechanism leverages mutual correlations between input components, making the model more physically interpretable compared to other architectures.

In our model, the input coordinates are first projected into a higher-dimensional embedding space through an embedding layer, followed by the addition of a learnable positional encoding to incorporate positional information. The resulting position-encoded representation is then passed through a stack of five transformer blocks, each consisting of a multi-head attention layer, a position-wise feed-forward network, residual connections, including layer addition and normalization. The output sequences from the final transformer block are aggregated using global average pooling across the sequence dimension, and the resulting pooled representation is finally passed through a dense layer to produce the estimated *Lp* of a given polymer. Detailed descriptions of the model implementation are provided in Appendix C 2.

#### 2. Data set construction and training procedure

Our DL machine is trained with a data set of simulated 2D WLC trajectories with constant persistence length. Mathematical details of generating simulation data are provided in Appendix A. As no specific profiles are assumed in advance, this approach not only facilitates the construction of the data set but also allows us to estimate any persistence length profiles without prior knowledge. More specifically, the *Lp* values ranging from 2 to 2 × 104 are uniformly grouped on a logarithmic scale, and 1.52 × 105 polymer trajectories are generated for each given *Lp*. We use 90% of the data set to update the model parameters (the training data), while the remaining 10% is reserved for the validation data.

Sampling data on a logarithmic scale enables the model to cover a wide range of *Lp*, and it improves its ability to distinguish between flexible and stiff polymers by providing distinct features in different stiffness regimes. Furthermore, training over a broad range of *Lp* facilitates the application of the model to diverse biological systems, as the persistence lengths of biopolymers span several orders of magnitude. For example, DNA has a persistence length of the order of tens of nanometers, actin filaments of the order of tens of micrometers, and microtubules up to several millimeters [13, 38, 39].

A primary quantity for the DL model is the logarithm of the persistence length, log_10_ *Lp*. This effectively scales the broad range of *Lp* in the training data and mitigates biased error contribution to the gradient, particularly for polymers with a large *Lp*. The mean absolute error (MAE) between log_10_ *Lp* and the output of the machine, denoted by *F****θ***, is used as the loss function for training:

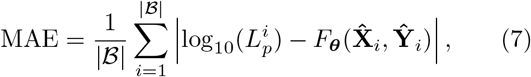

where |ℬ | is the batch size, 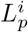 is the true persistence length of the *i*-th input polymer, ***θ*** is the model parameters, and 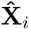 and 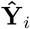 are the min-max scaled *x* and *y* coordinates of the *i*-th input polymer, respectively.

The model is optimized using the Adam optimizer with a learning rate of 10−4 and a batch size of 210, utilizing two NVIDIA RTX3090 GPUs. For efficient learning, the learning rate is reduced by a factor of 10−0.5 if the MAE loss for the validation data does not improve for 15 epochs. Additionally, the training is terminated if the validation loss does not improve for 50 epochs to prevent overfitting. The evaluation of the model on the validation data set is conducted at the end of each epoch. The model is implemented and trained using Keras with a Tensorflow 2.1 backend [40, 41].

All code materials used in this study, including the simulation of WLC conformations, data preprocessing, model architecture implementation, and model training, are available via our GitHub repository. See DATA AVAILABILITY STATEMENT for more information.

## III. RESULTS OF SIMULATED POLYMER CHAINS

To assess the accuracy and robustness of our DL method, we first consider a set of simulated polymer conformations. From simulations, we prepare polymers that have constant persistence lengths or position-dependent persistence lengths in a wide range of *Lp*. We also compare our DL prediction results with two conventional approaches based on the tangent-tangent correlation function and the end-to-end distance. Therefore, this provides a rigorous evaluation of our prediction model under controlled conditions where the ground truth of persistence length is known. In addition, we highlight the practical advantages of the proposed framework, particularly its robustness when only a limited number of polymer conformations are available.

### A. Polymers with a uniform bending persistence length

We evaluate the accuracy of our DL model for a set of conventional WLCs having a uniform bending modulus. For each value of *Lp* within the training range, 100 polymer samples of length *L* = 24 are generated using the simulation protocol described in Appendix A. The mean and standard deviation of *Lp* values inferred from our DL model are plotted as red circles in Fig. 2(a), compared to the ground truth (black dashed line). The result shows that our DL model produces accurate estimates of persistence lengths over a wide range of *Lp*.

**FIG. 2.**
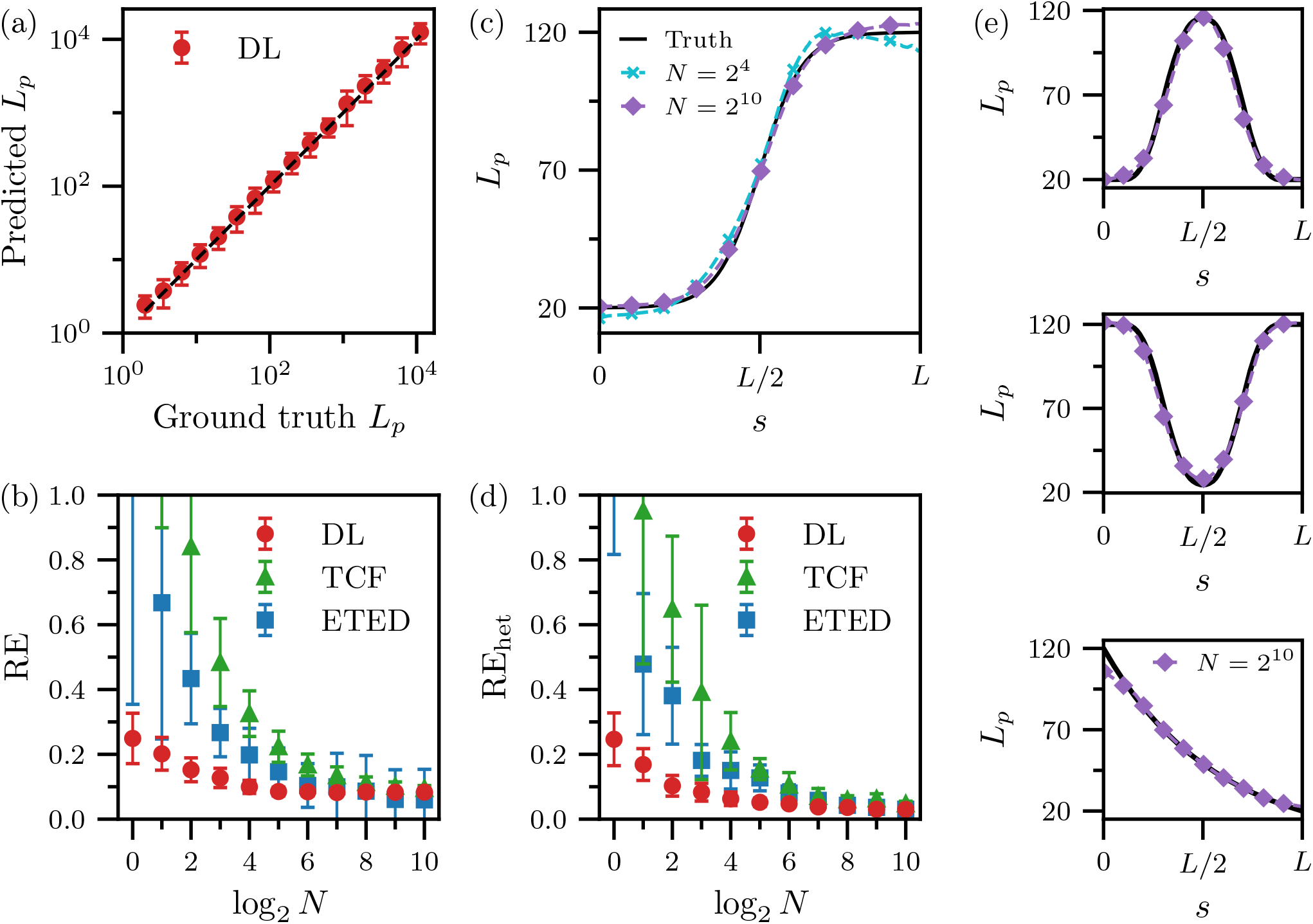
(a) Prediction of *L*_*p*_ from computer-generated conformations of WLCs having constant *L*_*p*_s using our deep-learning model (red circles), compared with the ground truth (black dashed line). (b) Relative Error of *L*_*p*_ from data sets of *N* polymer conformations, using the DL model (red circles), tangent–tangent correlation function (green triangles), and end-to-end distance method (blue squares). (c) Prediction of *L*_*p*_(*s*), a smoothly varying persistence length profile (black solid), using the DL method with *N* = 24 (cyan crosses) and *N* = 210 (purple diamonds). (d) RE of *L*_*p*_(*s*) from data sets of *N* polymer conformations, comparing the DL method (red circles), tangent–tangent correlation function (green triangles), and end-to-end distance method (blue squares). (e) Various profiles of *L*_*p*_(*s*) (black solid) and representative DL model predictions with *N* = 210 (purple diamonds).

We compare the performance of our DL model with the conventional methods using the tangent-tangent correlation function (TCF) and end-to-end distance (ETED), given in Eqs. (2) and (4), respectively. In the comparison, we investigate how the number of polymer samples, *N*, i.e, data availability, influences the prediction accuracy [Fig. 2(b)]. For a given *N*, a single persistence length estimate, denoted by *Lp,N*, is obtained, and the estimation accuracy is quantitatively measured by the relative error (RE):

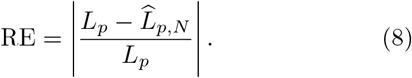

The RE metric ranges from 0 to + ∞, where it is equal to 0 when the estimation perfectly matches the true value, and increases as the estimation deviates further.

The mean and standard deviation of the RE(*N*), computed over 1.6 × 103 independent data sets for each *N*, are presented in Fig. 2(b). The DL model consistently produces more accurate predictions than the conventional ones in the entire range of *N* we used. In particular, while the conventional estimators exhibit substantial errors when *N* is small, the DL model maintains significantly low estimation errors under the same conditions. For example, at *N* = 2, the RE of our DL model is 7.2-fold and 3.3-fold smaller than those of the TCF and ETED, respectively. Furthermore, among the three methods, our model reaches the lowest RE most rapidly, converging only after *N* = 24. This clearly demonstrates the advanced efficiency of the DL model.

### B. Polymers with a spatially varying persistence length

Now we extend our approach to heterogeneous semi-flexible polymers characterized by a non-uniform persistence length along the contour. As a prototype example, we consider the semi-flexible polymers whose persistence length smoothly increases along the contour as plotted in Fig. 2(c). A data set of *N* such heterogeneous polymers of length *L* = 149 is generated using the protocol described in Appendix A. For these simulation data sets, we estimate the profile of persistence length, *Lp,N* (*s*), with our DL model and the protocol described in Appendix B. We examine the performance accuracy of our DL model in comparison with the ground truth profile [Fig. 2(c)]. For *N* = 24, the estimated *Lp* profile already closely follows the ground truth (cyan crosses). When the number of samples is increased to *N* = 210, the estimation becomes even more accurate (purple diamonds).

The prediction accuracy as a function of *N* is further quantified via the RE defined as

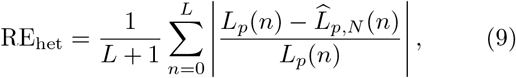

where *n* is the monomer index along the contour. The mean and standard deviation of the RE, computed over 10 independent data sets for each *N*, are shown in Fig. 2(d). Similarly found from the uniform *Lp* case, our DL method outperforms the conventional methods for the entire range of *N*, particularly when *N* is small.

In addition to the prototype profile in Fig. 2(c), we also investigate the performance of our DL method with several other profiles of *Lp*(*s*) shown in Fig. 2(e). Our DL method (symbols) accurately estimates the designed *Lp* profiles (solid lines) with *N* = 210 samples. In Fig. A3 (appendix), we provide the full analysis results for the four profiles in a wide range of *Lp* values in comparison with the conventional methods. For all profiles, it turns out that the estimation performances of the DL method exhibit a consistent trend as observed in Fig. 2(d). This robustness highlights the versatility of our DL method, which is trained with polymer chains with uniform persistence lengths.

We emphasize that a notable advantage of our DL method over the statistical methods is its ability to accurately predict persistence length profiles with a limited number of polymer data. For example, at *N* = 2, our method yields significantly lower estimation errors compared to the other methods, and as few as *N* = 22 samples are practically sufficient for reliable estimation. This capability is particularly relevant in the experimental context, because producing large data sets is often limited. Consequently, our method enables accurate prediction of *Lp* under such limited, real-world conditions.

## IV. APPLICATION TO BIOLOGICAL SYSTEMS

The above prediction of *Lp* based on the simulation data provides a rigorous foundation for evaluating the accuracy of our DL approach. Through quantitative comparisons with conventional methods for both uniform and spatially varying *Lp* of polymers, we have demonstrated that our proposed DL method consistently achieves superior performance. Based on this, now we extend to biological systems by analyzing experimental image data, which include B-DNA, actin filaments, filopodia, and intracellular microtubules. Our aim is not only to further validate the applicability of our DL method but also to elucidate structural features of complex biopolymers in terms of accurate persistence length prediction.

### A. B-DNA and actin filaments

B-DNA and actin filaments are biopolymers known to have persistence lengths of about 40–50 nm and 10– 20 *µ*m, respectively [38, 39]. For our DL prediction, we adopt experimental images of B-DNA obtained from atomic force microscopy [42], and fluorescence images of actin filaments in human mesenchymal stem cells [43], respectively. For the estimation, we extract 130 B-DNA chain conformations and 1007 actin filaments obtained using the tracking algorithms proposed in Refs. [31, 32]. Representative traced curves of B-DNA and actin filaments are shown in the top panels of Figs. 3(a) and (b), respectively.

**FIG. 3.**
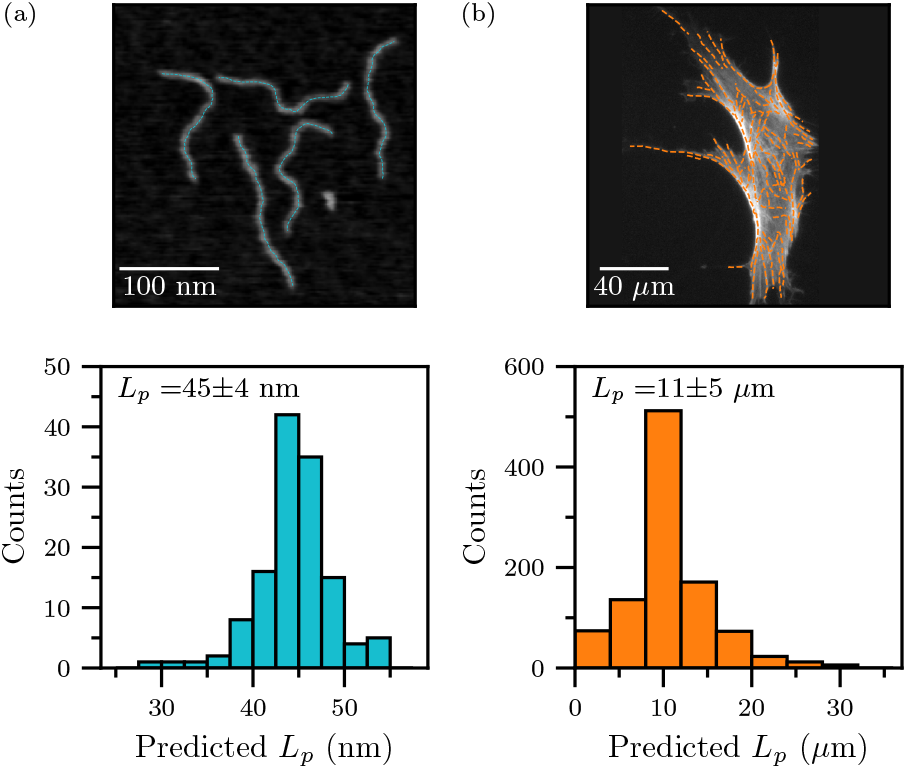
Top: Experimental images of (a) B-DNA and (b) actin filaments, with traced chain conformation curves overlaid as colored dashed lines. Below: Histograms of our DL model inferred *L*_*p*_ of the corresponding experimental images. The mean and standard deviation of *L*_*p*_ are written in each histogram.

The estimated *Lp* values of B-DNA and actin filaments through our DL model are plotted as histograms in the bottom panels of Figs. 3(a) and 3(b). As a mean value, we obtain *Lp* = 45 ± 4 nm for B-DNA and *Lp* = 11 ± 5 *µ*m for actin filaments, respectively, which are consistent with those reported in previous studies. In particular, the estimated range of *Lp* for B-DNA is in good agreement with the values (45 ± 3 nm) obtained using the ETED [7]. Summing up, these two examples demonstrate that our DL model generally works for experimental images of biopolymers with constant *Lp*s.

### B. Filopodia

Filopodia are cellular protrusions composed of tightly parallel-bundled F-actin filaments that extend from the plasma membrane [44]. The barbed ends of these filaments are oriented outward, allowing continuous actin polymerization at the tip, while their extension and retraction are regulated by retrograde flow and adhesion dynamics [45]. Functionally, filopodia are critical for sensing the extracellular environment, directing cell migration, and facilitating adhesion. These structures are equipped with surface receptors that detect external chemical cues and trigger downstream intracellular signaling pathways [46].

Investigating the mechanical properties of filopodia is challenging, as each individual filopodium interacts differently with actin-binding proteins and the surrounding environment, leading to diverse structural features in length, bending stiffness, and thickness. Only a limited number of studies have reported the persistence length of filopodia. For example, Zidovska *et al*. reported *Lp* ≈ 1 mm for in vivo filopodia by analyzing elongation using magnetic tweezers [47]. Chang *et al*. estimated *Lp* = 957 *µ*m–1.39 mm for filopodia in live HeLa cells by force-extension experiments and molecular dynamics simulations [45]. These studies rely on controlled experimental setups to precisely measure the responses of filopodia against external forces, followed by theoretical frameworks to interpret the responses, which is often challenging due to the complex and heterogeneous nature of the cellular environment.

In contrast, several studies have employed a bundle of actin filaments in vitro as a simplified system of filopodia. Claessens *et al*. reported that actin-binding proteins strongly influence the bending stiffness of F-actin bundles in vitro [48]. Takatsuki *et al*. demonstrated that the persistence length of F-actin bundles ranges from 10 *µ*m (the value for a single filament) up to 150 *µ*m as the level of fascin (a crosslinking protein of F-actin filaments) increases in vitro [49]. This simplified system has been adopted in various filopodia studies but cannot fully describe live-cell systems, since filopodia in living cells are enclosed by the plasma membrane and sensitively influenced by actin-binding proteins and other intracellular molecules [47, 48, 50–52]. Given these complexities, the persistence length of a filopodium in living cells is expected to be spatially varying along its contour.

In this context, we apply our DL model to overcome these limitations. For this, we analyze fluorescence microscopy images of filopodia in HeLa cells. To visualize F-actin filaments within filopodia, we label them using Lifeact, a 17-amino acid peptide that specifically binds to F-actin. Live-cell imaging is performed using highly inclined and laminated optical sheet microscopy. Detailed descriptions on the imaging method are provided in Appendix E. From the fluorescence images, we extract six distinct filopodia morphologies from tip to base, using the same tracking algorithm described in the previous section. The extracted morphologies are shown in Fig. 4(a). Subsequently, we apply our method to predict position-dependent persistence length profiles of each filopodium.

**FIG. 4.**
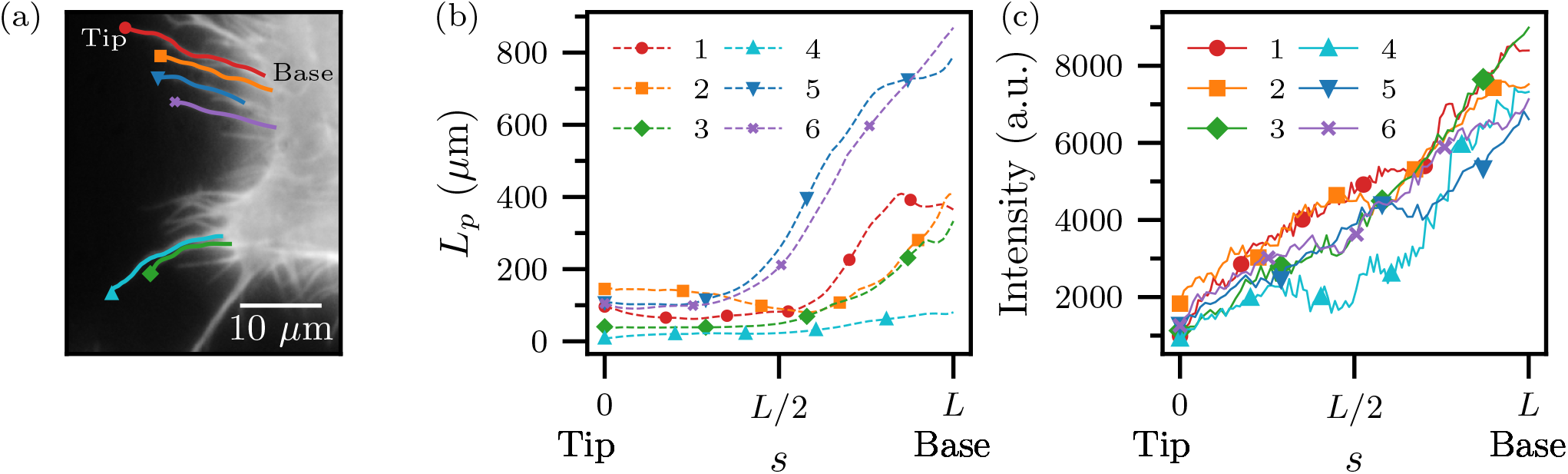
(a) Traced morphology curves of six individual filopodia from tip to base. (b) Persistence length profiles of each filopodium. The arc length coordinate *s* = 0 corresponds to the tip, and *s* = *L* to the base. (c) Fluorescence intensity profiles along the contour of each filopodium.

The persistence length profiles estimated for all six filopodia are plotted in Fig. 4(b). We note that the estimated values of *Lp* range from several tens of micrometers to nearly a millimeter. This is consistent with the previously reported measurements in live-cell filopodia studies, i.e., *Lp* = 1 mm in Ref. [47] and *Lp* = 957 *µ*m– 1.39 mm in Ref. [45], respectively. Another important feature of the filopodia’s bending stiffness is that the *Lp* tends to increase from the tip to the base. This morphological pattern is in line with the underlying biological mechanisms of filopodia protrusion. The extension of filopodia is driven by F-actin polymerization, which is implemented by the diffusion of G-actin monomers within the filopodia shafts [53]. This process typically results in a tapered morphology at the tip compared to the base, leading to smaller bending stiffness near the tip compared to the base. The fluorescence intensity profiles shown in Fig. 4(c) further support this observation. For all cases, the intensity increases from the tip to the base, reflecting increases in the thickness of the filopodial structure.

The substantial variation in persistence length along the filopodia’s contour can be further interpreted in the context of F-actin bundle mechanics. Experimental studies have reported that the stiffness of F-actin bundles depends on the number of filaments *N* and their crosslinking density [47, 48, 50–52, 54, 55]. The persistence length of a bundle is modeled as *Lp* ≈ *I*(*N*)*Lp*,single, where *Lp*,single ~ 10 *µ*m is the persistence length of a single Factin, and *I*(*N*) is a scaling factor. It has been reported that actin-binding proteins sensitively determine F-actin bundle stiffness; *I*(*N*) ~ *N* for loosely bundled filaments and *I*(*N*) ~ *N* 2 for tightly bundled ones. Adopting a bundle size of *N*≈ 7 from Ref. [45], we roughly estimate that the persistence length of filopodia ranges from about 70 *µ*m (weak bundling) to over 500 *µ*m (tight bundling). These estimates are in good agreement with the values inferred from our DL-based profiles. From these results, we speculate that the distal, newly polymerized regions of filopodia remain mechanically soft, while the proximal base becomes more rigid through tight bundling.

### C. Microtubules in living cells

Microtubules are fundamental components of the cytoskeleton in eukaryotic cells, playing crucial roles not only in maintaining cellular structure but also in facilitating diverse intracellular processes. The repeating units of *α*- and *β*-tubulin polymerize microtubules to form a hollow cylindrical structure. This highly ordered structure results in significant mechanical strength with inherent resistance to both bending and compression, allowing it to support the cytoplasmic organization [56]. Microtubule polymerization occurs in a head-to-tail manner, producing polarized filaments characterized by a fast-growing plus end and a slow-growing minus end. This intrinsic polarity provides directionality in intracellular transport mediated by motor proteins such as kinesin and dynein. In addition to transport, microtubules play a critical role in physically segregating chromosomes and orienting the cleavage plane during cell division, and also serve as the principal structural element of motile organelles such as flagella and cilia [57].

Numerous studies have characterized the structural, mechanical, and dynamical properties of micro-tubules [13, 58, 59]. Several investigations have revealed that microtubules behave differently within living cells. Pampaloni *et al*. experimentally and theoretically demonstrated that the persistence length of microtubules depends on their total contour length, attributed to the substantial disparity between the longitudinal Young’s modulus and the shear modulus [60]. It was also reported that the persistence length of microtubules in live cells is approximately 20–30 *µ*m, significantly shorter than the values measured in vitro [61, 62]. This reduction is primarily driven by non-thermal forces in cells, particularly those generated by ATP-consuming motor proteins. Moreover, various microtubule-associated proteins influence microtubule mechanics, and post-translational modifications, such as acetylation and deacetylation of microtubules, were shown to modulate the bending stiffness, which often occur heterogeneously in individual microtubules [12, 63–66]. Consequently, the elastic properties of microtubules in living cells are expected to be more spatially heterogeneous compared to in vitro observations.

To apply our DL method, we analyze fluorescence microscopy images of microtubules in HeLa cells. Microtubules are visualized using a fluorescent probe that specifically labels them, and live-cell imaging is conducted using Airyscan microscopy. Detailed descriptions of the imaging technique are provided in Appendix F. Representative fluorescence images of HeLa cells are shown in the top panels of Figs. 5(a) and 5(b). From these images, we obtain morphological curves of the microtubules using the same tracking algorithm described previously, and their persistence lengths are subsequently predicted using our DL method.

**FIG. 5.**
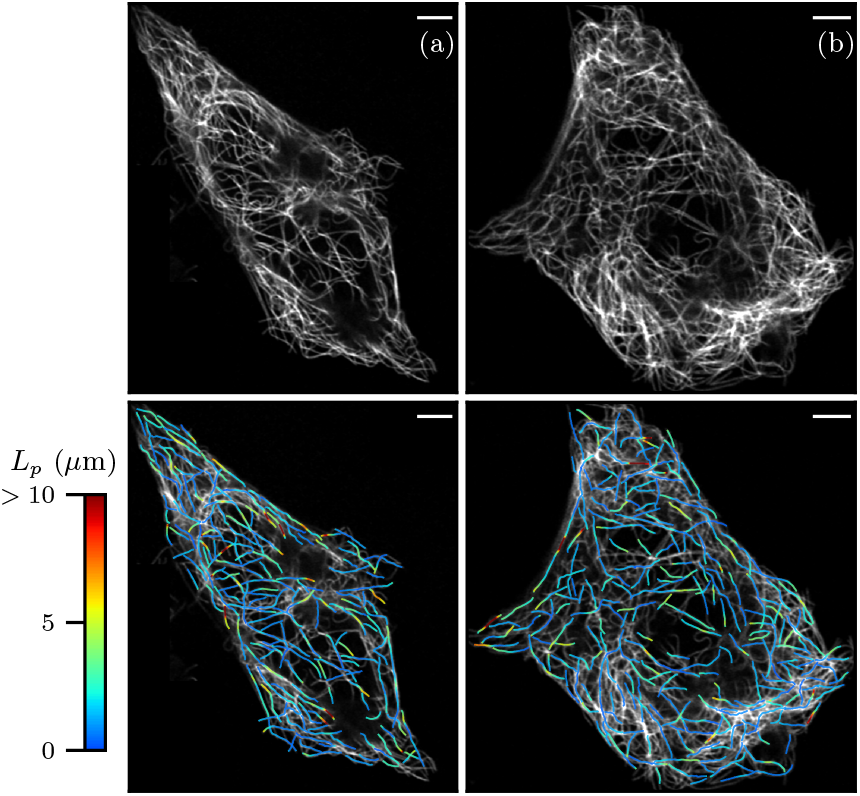
Top: Experimental fluorescence images of two distinct HeLa cells [(a) and (b)] in which microtubule structures are visualized. Bottom: Traced morphology curves of microtubules colored according to the predicted persistence lengths. Values of *L*_*p*_ greater than 10 *µ*m are depicted by red color. The scale bar depicts 5 *µ*m.

In the bottom panels of Figs. 5(a) and 5(b), we present the traced microtubule morphological curves with distinct colors according to the values of persistence lengths. We observe that microtubules in two distinct cells display substantial variability in persistence length along their contour. As discussed previously, such heterogeneities are expected because of differences in the length of microtubules and their interactions with a variety of intracellular components. The estimated persistence lengths span a range from a few micrometers to approximately 30 *µ*m, consistent with the values reported in the previous live-cell measurements [60–62]. In particular, most values are smaller than 10 *µ*m, which may reflect enhanced flexibility near the cell periphery, where microtubules interact with the cell membrane and cortex frequently.

## V. DISCUSSION

In recent years, significant breakthroughs in machine-learning techniques [35, 67, 68] have led to notable applications across a wide range of disciplines [21, 36, 37, 69]. Despite these successes, one of the major challenges for machine-learning models is the so-called ‘black box’ problem, where the internal mechanisms of the algorithms remain opaque. To overcome this, two main approaches have generally been suggested. The first one incorporates prior physical knowledge directly into the learning process by embedding known physical laws or constraints in the model architecture or loss function, thereby enhancing both interpretability and scientific relevance [19, 70, 71]. The second one investigates how different parts of input data contribute to output by directly analyzing model’s internal computation processes [72– 74]. In this section, we integrate both perspectives and further analyze to elucidate the inner workings of the machine used in our study.

### A. Relationship with physical quantities

To quantitatively analyze the features leveraged by our DL model, we investigate the latent space of the global average pooling layer, where the outputs of all transformer blocks are aggregated (see Fig. A2 for the model architecture). Using 1.6 × 104 polymer chains of uniform persistence lengths and *L* = 24, we extract 64-dimensional feature vectors, respectively, corresponding to a single polymer chain conformation. We then train an XGBoost regressor model to estimate *Lp* from these feature vectors and obtain the normalized feature importance against the 64 features, as shown in Fig. 6(a) [75]. It turns out that the 64 features are not equally important for the estimation of *Lp*. There are only a few features that play a significant role in inferring the persistence length (e.g., the feature of index 36 is the most dominant one).

**FIG. 6.**
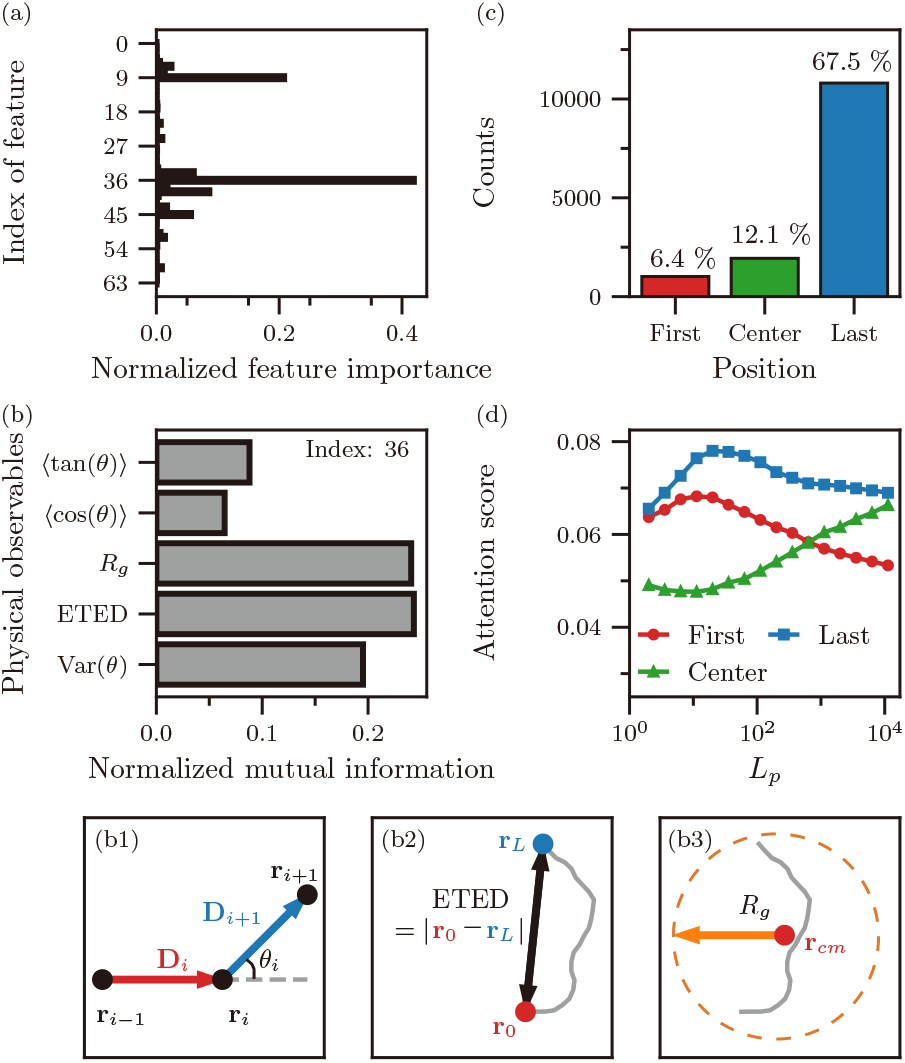
(a) Normalized feature importance vs. index of feature from the global average pooling layer, identified by an XGBoost model. The feature importance of a feature *f*_*i*_ is calculated according to the definition ∑ _*t ∈ T n*_ ∑_*N ∈t*_ 1*f*(*n*)=*f*_*i*_, where *T* is the set of all decision trees in the XGBoost model, *N*_*t*_ the number of nodes in the tree *t*, and 1 is an indicator function that equals to 1 if *f*_*i*_ is used at node *n* (i.e., *f* (*n*)), and 0 otherwise. The feature importances are normalized for their sum to be unity. (b) Normalized mutual information between the most important latent feature (index 36), associated with various physical quantities–(b1) The variance of turning angles (Var(*θ*)), (b2) the absolute value of end-to-end distance (ETED), and (b3) radius of gyration (*R*_*g*_). (c) The number of polymer samples whose maximum self-attention score occurs at the first (red), center (green), and last (blue) diagonal position groups. Percentage values shown above each bar represent the fraction of each position group. (d) Mean self-attention scores at the first (red circles), center (green triangles), and last (blue squares) groups, plotted as a function of *L*_*p*_.

Given the feature importance, we further calculate five physical observables for each polymer chain against the feature vectors. They are: the variance of turning angles (Var(*θ*)), the absolute value of end-to-end distance (ETED), radius of gyration (*Rg*), mean cosine of the turning angles (⟨cos(*θ*) ⟩), and mean tangent of the turning angles (⟨ tan(*θ*) ⟩). See the schematics in Fig. 6(b1)– (b3) for visual definitions of these quantities. Mathematical definitions of the five observables are explained in Appendix G 1. Subsequently, we quantify the correlation between the most dominant five latent features and the above physical observables using the normalized mutual information (NMI), given as

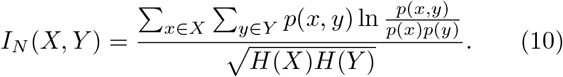

Here, *X* and *Y* are random variables, *p*(*x, y*) denotes their joint probability distribution, *p*(*x*) represents the marginal probability of *X*, and *H*(*X*) = −∑_*x ∈ X*_ *p*(*x*) ln *p*(*x*) corresponds to the Shanon entropy of *X* [76]. The NMI ranges from 0 to 1 and quantifies the statistical dependence between *X* and *Y*. The NMI is 0 when *X* and *Y* are independent, while higher NMI values reflect stronger correlations between them.

In Fig. 6(b), we present the measured NMI between the most dominant latent feature (index of 36) and the five physical observables. The corresponding NMI values for the remaining four latent features can be found in Fig. A4. For the five dominant latent features, Var(*θ*), ETED, and *Rg* unanimously exhibit higher NMI values compared to the other quantities. While Var(*θ*) captures the information of local bending fluctuations, ETED and *Rg* measure the global geometric feature of a polymer chain conformation. This suggests that our deep-learning model sees both global and local geometric features of a chain’s bending configuration when inferring the persistence length.

### B. Self-attention scores

We further analyze the self-attention matrices to investigate how the relevant physical quantities discussed earlier are related to the internal mechanism of our DL model. Visualizing these matrices is a common approach for understanding a transformer-based deep-learning model, as it reveals how the model distributes attention across different elements of the input sequence. This analysis provides insights into which regions of the input the model is focusing on when making predictions [35]. Using the same polymer conformation set used in the previous section, we compute the self-attention matrices for all attention heads of five transformer blocks. These matrices are then aggregated into a single matrix, referred to as the averaged self-attention matrix, as described in Appendix G 3. We then visualize the resulting matrices as heatmaps in Fig. A5. We observe that all averaged self-attention matrices exhibit relatively higher self-attention scores in the diagonal positions and its nearest neighbor regions across the entire range of *Lp*. These patterns suggest that the model captures local structural features, such as turning angles between adjacent monomers.

As shown in Fig. 6(c), we examine the underlying patterns of the averaged self-attention matrices (see Fig. A5). For a single averaged self-attention matrix, denoted by *Ā*, we identify the entry exhibiting the highest self-attention score and categorize it into one of three groups depending on the diagonal position: the first (*Ā*_1,1_), central (*Ā*_13,13_), or last (*Ā*_25,25_). Here, *Ā*_*i,j*_ denotes the (*i, j*)-th entry of the averaged self-attention matrix with *i, j* ∈ [1, 25]. The results show that approximately 86 % of the highest self-attention scores in the averaged self-attention matrices are concentrated at either the first, central, or last diagonal position. Moreover, the first and last positions consistently exhibit pronounced vertical lines across the entire range of *Lp*, indicating that the model treats these two endpoints as reference points (see Fig. A5). This behavior can be attributed to the model’s encoding of global geometric features, such as the end-to-end distance. This observation also reflects the preprocessing procedure. As described in Appendix C 1, the initial orientation of a polymer chain is aligned along the positive *x*-axis prior to input. Under this transformation, the position of the last monomer in a flexible polymer tends to deviate more significantly from the *x*-axis, whereas in a stiff polymer it remains relatively close. This structural variation likely leads to the higher self-attention scores associated with the first and last positions.

In contrast, the higher scores in the central group are relevant to the radius of gyration. This is because *Rg* quantifies the spatial distribution of monomers with respect to the center of mass, which is typically located near the central monomer along the polymer chain, particularly in stiffer polymers. Figure A5 further supports this idea. The averaged self-attention matrices begin to exhibit significant vertical lines at the central group if *Lp* ≳ 2 × 101. As discussed earlier, these vertical lines indicate that all matrix elements concentrate on the center, suggesting that the model collectively attends to the central monomer as a pivotal point for extracting global shape descriptors, which are likely associated with the radius of gyration.

We further investigate how the self-attention scores of the first, central, and last groups vary with respect to *Lp*. As shown in Fig. 6(d), the results exhibit non-monotonic trends for all three groups. The scores corresponding to the central group reach a minimum near *Lp* ~ 2 × 101 and gradually increase with increasing *Lp*, consistent with the previous observation. In contrast, the scores for the first and last groups peak around *Lp* ~ 2 × 101, and subsequently decrease. Notably, the last group consistently exhibits higher self-attention scores than the first and central groups across the entire range of *Lp*, possibly reflecting its geometric importance in defining global features and an important role during the preprocessing. Additionally, we note that the self-attention scores for the first, central, and last groups all exhibit a change in trend near *Lp* = 2 × 101, which is notably close to *Lw* = 24. This observation suggests that the model may adjust its attention strategy as *Lp* approaches the contour length of the input polymer, shifting its focus toward global measures, such as the radius of gyration. This adaptive strategy may contribute to the model’s capacity to accurately estimate a wide range of *Lp*, significantly exceeding the window size *Lw*.

## VI. CONCLUSIONS

We have developed a deep-learning framework that uses two-dimensional coordinates data as input to comprehensively probe the elastic properties of biopolymers in terms of a spatially varying persistence length. Our DL method consists of three core steps: (i) segmentation of polymer chain conformations into overlapping segments, (ii) pre-estimation applying a transformer-based neural network to infer the persistence length of each segment, and (iii) reconstruction of the full persistence length profile using the sliding-window average. The model was trained on a data set of simulated two-dimensional position data from semiflexible polymer conformations with uniform persistence lengths. The resulting framework outperformed the conventional methods (based on the tangential vector correlation and the end-to-end distance) in estimating persistence lengths, both uniform and even position-dependent. Our DL model achieved high estimation accuracy in a wide range of persistence length profiles, which remains robust even with a limited number of available samples. This prediction performance under limited data availability significantly exceeded that of the conventional approaches requiring large data sets. This represents a key advantage of our model for experimental applications where data is often limited.

Given the trained framework, we further validated our DL model on various biopolymer systems. Using experimental images of B-DNA and actin filaments, our model predicted the average persistence lengths of 45 ± 4 nm and 11 ± 5 *µ*m, respectively, consistent with previously reported values. Furthermore, we demonstrated the capability of our method by applying it to more complex biological systems. Using our predictive model, we found that the persistence length of filopodia gradually increases from tip to base, consistent with their structural polarity and dynamic assembly mechanisms. We also applied our model to intracellular microtubules in living cells. The results revealed spatially heterogeneous persistence lengths under live-cell conditions, reflecting elastic and mechanical diversities induced by intracellular forces, molecular crowding, and the influence of microtubule-associated proteins.

Consequently, we demonstrated that our predictive model outperforms the conventional approaches and efficiently captures elastic features of biopolymers in diverse cellular systems. Moreover, we discussed and provided deeper insights into how and why the deep-learning model achieves its superior performance. By analyzing latent feature representations and self-attention patterns, we showed that the model leverages multiple physically relevant quantities, particularly the variance of turning angles, end-to-end distance, and radius of gyration, which are interpretable and closely related to polymer stiffness. Notably, the model adapts its attention strategy in response to the stiffness of the input polymers. These findings not only enhance the interpretability of the model but also establish a connection between deep-learning and polymer physics.

To conclude, we have developed a robust, accurate, and interpretable deep-learning framework, which predicts a persistence length of polymers from their chain conformation data alone. This framework is broadly applicable to investigating the elastic properties of biopolymers, of which knowledge is essential for studying the mechanical effects induced by diverse interactions inside cells. Our predictive model, fine-tuned in both its methodological strategies and deep-learning architecture, may serve as a standard protocol to inspire future research in data-driven modeling of complex biological and soft matter systems.

## ACKNOWLEDGMENTS

This work was supported by the National Research Foundation (NRF) of Korea, Grant No. RS-2023-00280169, RS-2023-00260454, RS-2023-00218927, & RS-2024-00343900. The authors thank Jaesung Choi for insightful discussions regarding the discussion section.

## DATA AVAILABILITY STATEMENT

All code materials used in this study are available on the following GitHub repository: https://github.com/ChangbeomHong/Lp. This repository includes scripts for simulating polymer chain conformations of the worm-like chain model with both uniform and spatially varying persistence lengths based on the discrete worm-like chain model, as well as for implementing the deep-learning model used in this study. Additional data supporting the findings of this study are available from the corresponding author upon reasonable request.

## Appendix A

### Computer-generated conformation of polymer chains

We present details of the discrete two-dimensional worm-like chain (WLC) model and our simulation protocol for generating multiple equilibrium WLC conformations. In units of bond length *a* ≡ 1, the chain comprises *L* bonds and *L* + 1 monomers whose positions are denoted by 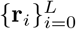. Here, the monomer index *i* also serves as the discrete arc-length parameter. The discrete version of the bending energy (effective Hamiltonian), given in Eq. (5), is

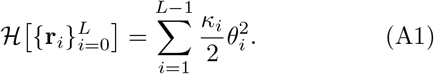

Here, *κi* is the bending modulus of the *i*-th monomer, and *θi* is the angle between the successive bond vector **D***i* and **D***i*−1, where **D***i* ≡ **r***i* − **r***i*−1. The local persistence length of the *i*-th monomer in two dimensions (2D) is

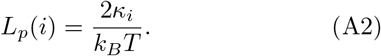

Therefore, the Boltzmann distribution leads to

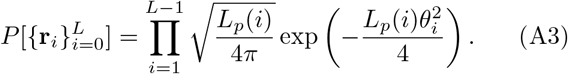

Here, *θi* is an independent and identically distributed random variable following the normal distribution of zero mean and variance 2*/Lp*(*i*). The chain has a constant persistence length in the limiting condition, *Lp*(*i*) = *Lp* for all *i*.

To generate a chain of length *L*, we first sample the {*θi*} from the Boltzmann distribution (A3). The polymer’s monomer position is then constructed via

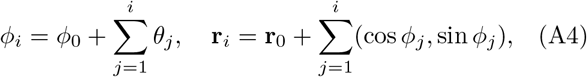

where *ϕi* is the polar angle of the **D**_*i*+1_ with respect to *x*-axis. Without loss of generality, we set **r**_0_ = (0, 0) and *ϕ* _0_ = 0. We repeat this process to generate a set of polymer conformation.

## APPENDIX B

### Reconstruction procedure of persistence profile lengths

Figure A1(a) shows the details of segmentation step, which is the first process in our DL method. We begin with a subset consisting of generated *N* polymers of length *L* (in a black dashed box). Each polymer conformation is divided into *L*− *Lw* +1 overlapping segments of a sliding window of length *Lw*, producing *N* segments at each window position (black solid boxes). For each group of *N* segments, the DL model is applied to estimate *Lp* values. At this stage, we also use conventional methods to estimate *Lp* by computing the tangent-tangent correlation function [Eq. (2)] and the end-to-end distance [Eq. (4)] from all *N* segments.

As shown in Fig. A1(b), the pre-estimation step produces a sequence of estimated *Lp* values, denoted by 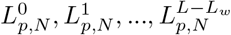. The final persistence length profile is then reconstructed via the sliding window averaging,

**FIG. A1.**
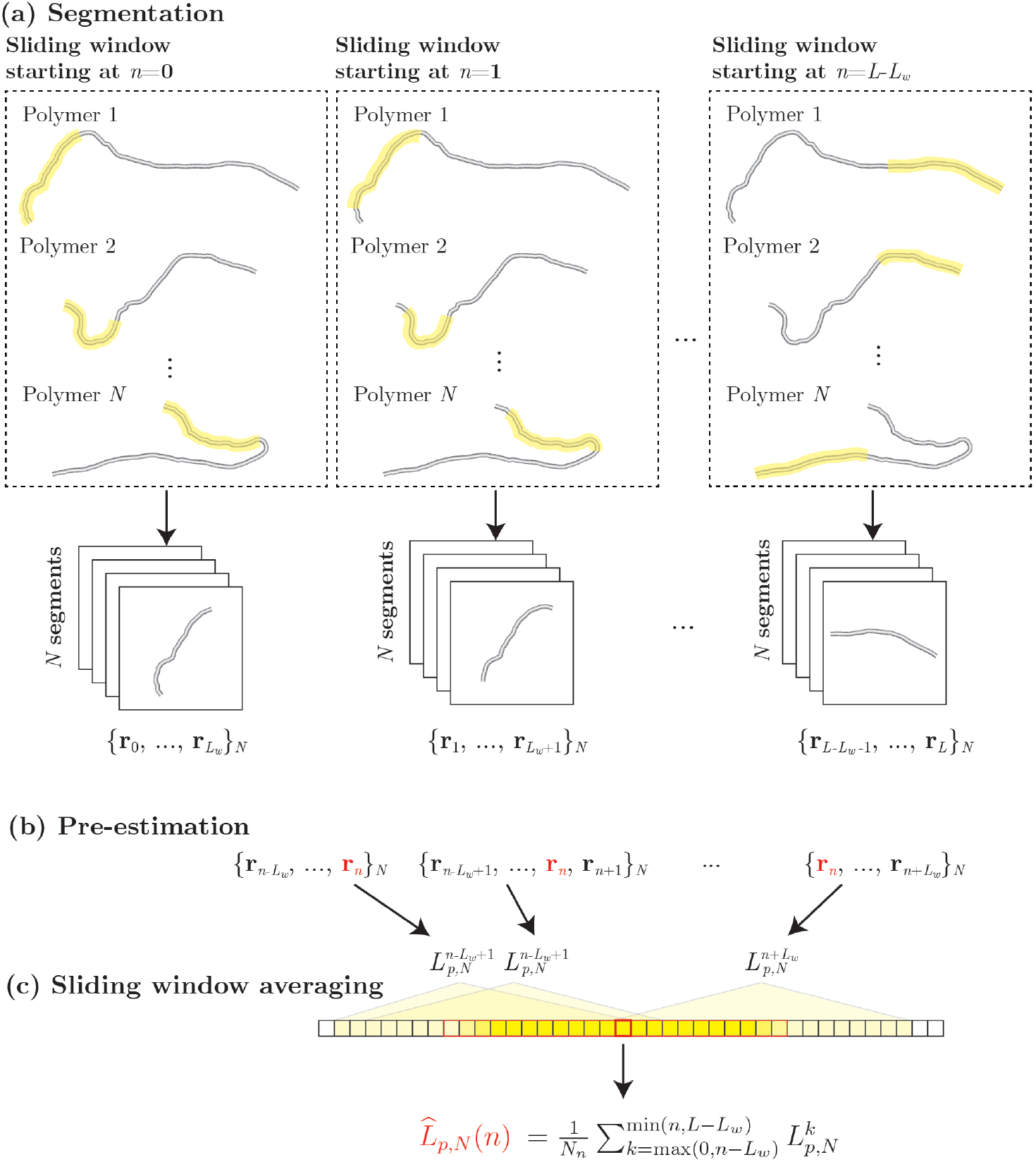
(a) Segmentation: For a subset of *N* polymer chain conformations (black dashed box), each polymer conformation is divided into *L − L*_*w*_ + 1 overlapping segments, resulting in *N* segments at each window position *n* (black solid boxes). (b) Pre-estimation: For each group of *N* segments extracted by a window at position *n* = *i*, the persistence length, denoted by 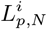, is estimated using our DL model, conventional tangential vector correlations, and end-to-end distance relations. (c) The persistence length profile is reconstructed by averaging the set of estimated value 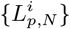.

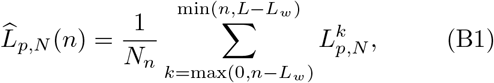

where 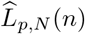 is the averaged persistence length at *n*-th monomer index, *n* ∈ [0, *L*], and *Nn* = min(*n, L* − *Lw*) − max(0, *n* − *Lw*) + 1 is the number of segments that the monomer of index *n* is included [see Fig. A1(c)]. Namely, the persistence length at a given monomer is computed as the mean of all local persistence lengths for the segments that include the monomer.

## APPENDIX C

### Data preprocessing and model architecture

#### 1. Preprocessing of polymer conformation data

We present details of the preprocessing procedures for WLCs of length *Lw* = 24, consisting of *Lw* + 1 = 25 monomers with position vectors denoted as 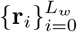. As a first step, each polymer is rotated such that its end-to-end vector is aligned along the direction of *x*-axis, yielding a new position vectors 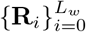. From the latter, we extract two coordinate vectors: 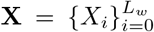 and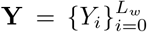, which respectively contain *x* and *y* coordinates of all monomers. Each of these vectors is independently scaled using a min-max scaler:

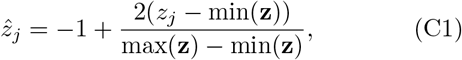

where 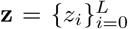 represents an input vector (either **X** or **Y**), and 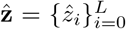 is the corresponding scaled vector (either 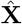 or **Ŷ**). This scaler transforms an input vector **z** such that the minimum and maximum values of the resulting vector 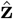 is −1 and +1, respectively. The scaled vectors 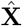 and **Ŷ** are used for the two input channels for the DL model, represented as a two-channel one-dimensional array of length *Lw* + 1.

#### 2. Deep-learning model architecture

As illustrated in Fig. A2, our DL model is built upon the transformer encoder architecture [35]. A two-channel one-dimensional array of length 25, consisting of the minmax scaled x and y monomer coordinates of a polymer, is first embedded into a higher-dimensional embedding space with *d*embed = 64 using a fully connected layer (Embedding layer). To incorporate positional information, a trainable positional embedding is then added to the embedded representation (Positional embedding).

**FIG. A2.**
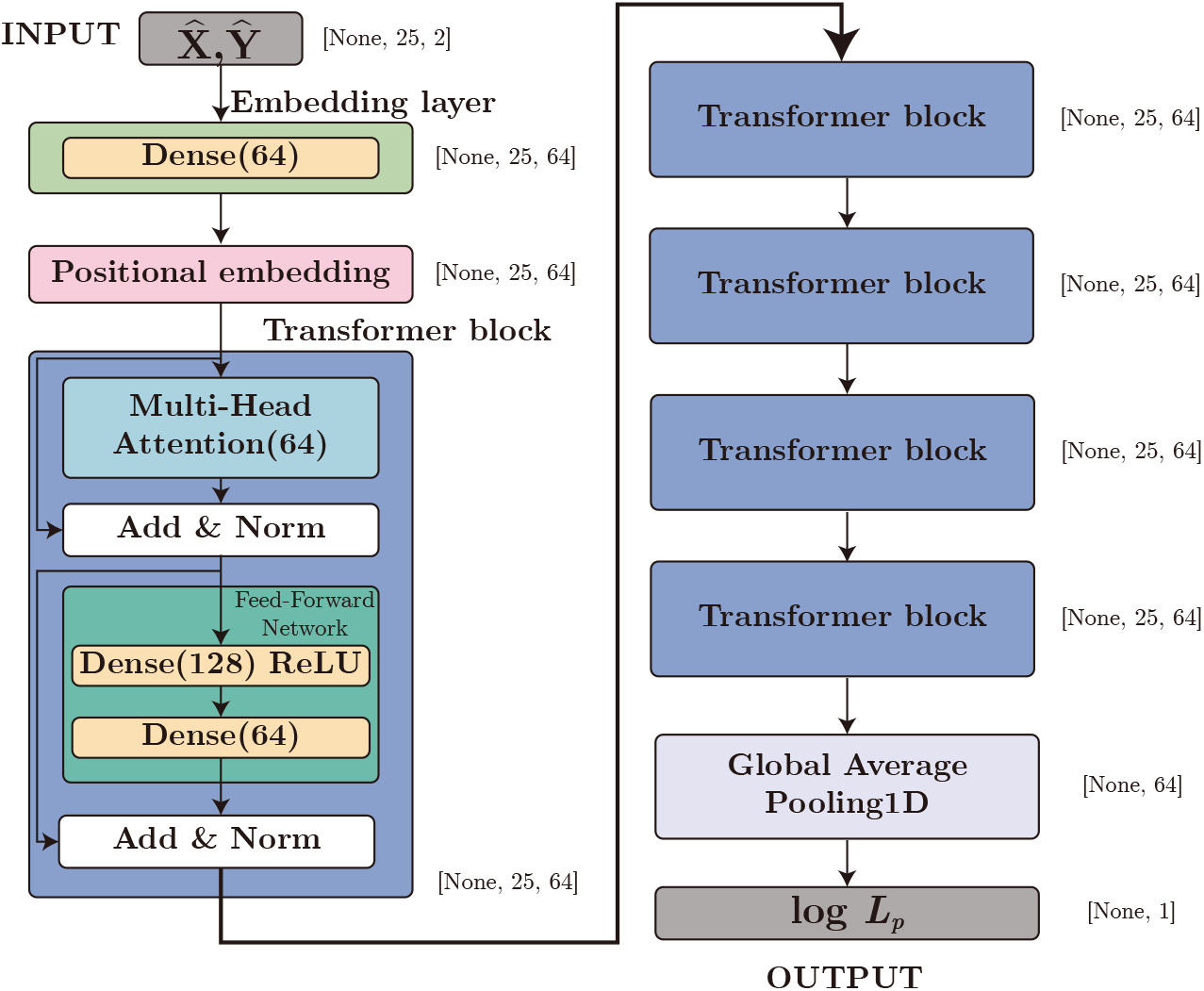
Architecture of our deep-learning model. Black arrows indicate the data flow from input to output. Key components (embedding layer, positional embedding layer, transformer block, and global average pooling layer) are shown with detailed specifications. Within each box, the number of perceptrons in fully connected layer (Dense) and the number of heads in multi-head attention layer are shown in parentheses. ‘ReLU’ denotes the use of the ReLU activation function. Output shapes are specified in brackets next to each layer or module, where ‘None’ denotes a variable dimension depending on the batch size.

Subsequently, a stack of five transformer blocks is consecutively applied to extract relevant features for persistence length estimation from the position-encoded representation. Each transformer block primarily consists of three components: a multi-head self-attention layer, layer addition and normalization with residual connections, and a position-wise feed-forward network. The multi-head attention layer learns positional dependencies from the input representation using 64 attention heads (Multi-Head Attention), after which the output is combined with the input through a residual connection followed by normalization (Add & Norm). The normalized representation is fed into the position-wise feed-forward network consisting of two dense layers, which expands the attention outputs to a higher-dimensional space of size *d*FFN = 128 before projected back to *d*embed. After application of the feed-forward network, another residual connection and normalization are applied to refine the representation.

The output representation from the last transformer block is aggregated using a global average pooling layer across the sequence dimension (Global Average Pooling 1D). The pooled features are then used to estimate the logarithm of the persistence length of the input polymer.

## APPENDIX D

### Estimation results for various heterogeneous profiles

We further benchmark the performance of our method by applying to additional profiles that are not presented in Section III B. As shown in Fig. A3, we consider four distinct profiles (a–d) across three different ranges of *Lp* (1–3). Following the same procedure previously described, we compute the RE of the predicted persistence length profiles using all three methods for different *N*. For each *N*, we present the mean and standard deviation of the RE, calculated over 10 independent test data sets.

**FIG. A3.**
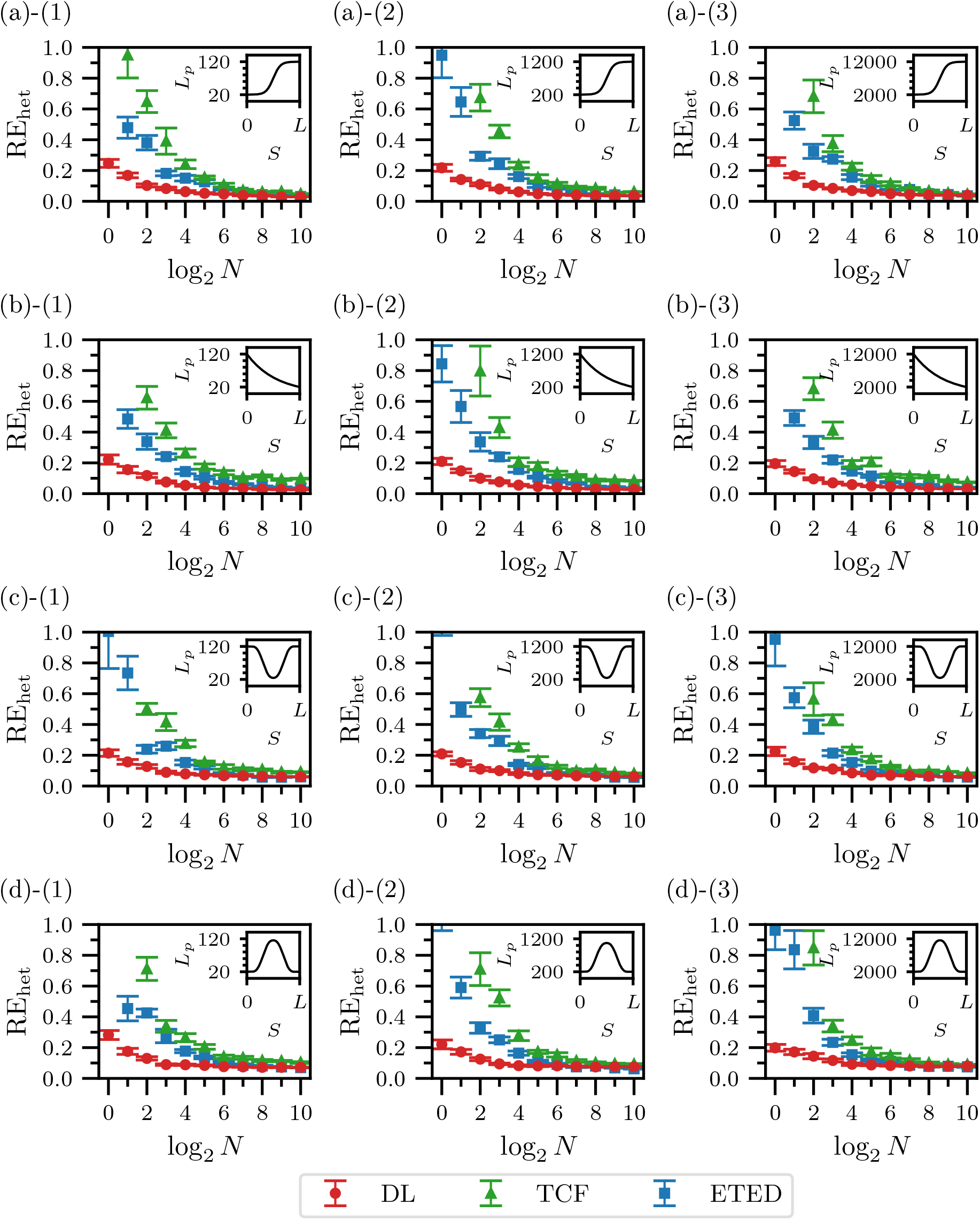
RE of predicted persistence length profile from data sets of *N* polymer conformations, comparing the deep-learning method (red circles), tangent-tangent correlation function (green triangles), and end-to-end distance method (blue squares). The analysis considers four smoothly varying profiles (a–d), each evaluated across three scaling ranges of *L*_*p*_: (1) 20–120, (2) 200–1200, and (3) 2000–12000.

## APPENDIX E

### Experimental procedure for fluorescence imaging of filopodia

To acquire the fluorescence images of filopodia shown in Fig. 4, HeLa cells (Korean Cell Line Bank) were cultured in phenol red-free Dulbecco’s Modified Eagle Medium (DMEM) supplemented with 10% fetal bovine serum, 1× penicillin-streptomycin (Thermo Fisher Scientific), 1× GlutaMAX (Thermo Fisher Scientific), and 1× MycoZap (Takara). F-actin filaments in filopodia were labeled with Lifeact-EGFP, a 17-amino acid peptide that specifically binds to F-actin. Live-cell imaging was performed using highly inclined and laminated optical sheet (HILO) microscopy with a 100× objective lens (Olympus UPlanSApo 100×, NA = 1.4). Fluorescence signals excited by a 488 nm laser (Cobalt) were detected using an electron-multiplying charge-coupled device (EMCCD) camera (Andor iXon 897). Image acquisition and processing were carried out using MetaMorph software.

## APPENDIX F

### Experimental procedure for fluorescence imaging of microtubules in HeLa cells

To obtain the fluorescence images of microtubules shown in Fig. 5, HeLa (ATCC, CCL-2) cells are seeded 35 mm Glass bottom dish with 20 mm micro-well #1 cover glass (Cellvis, D35-20-1-N) and grown in Dulbecco’s modified Eagle’s medium (DMEM; Serena, MCL-002-500ml) supplemented with 10% fetal bovine serum (FBS; Welgene, S001-01) under 37*°*C, 5% CO2, for 24 hours. Cells are washed with pre-warmed imaging media (HBSS, calcium, magnesium, no phenol red, Gibco, 14025-092, supplemented with 10% FBS) three times and then stained with 1 mL imaging media containing 1 *µ*M Tubulin TrackerTM Deep Red (InvitrogenTM, T34077) under 37*°*C, 5% CO2 for 45 min. Cells are washed with pre-warmed 1 mL of imaging media 3 times again, and plate is filled with 1 mL of pre-warmed imaging media. Live-cell Images are acquired with 63x objective of Zeiss LSM900 with Airyscan 2 microscope and undergone airyscan processing (Zeiss, Zen 3.10 software).

## APPENDIX G

### Relevant physical quantities and averaged self-attention matrices

#### 1. Definition of physical observables

For a single polymer of length *L*, consisting of *L* + 1 monomers located at position 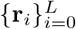, we calculate the following physical quantities.

##### a. Variance of turning angles

For the displacement vector **D***i* = **r***i* − **r***i*−1, the turning angle *θi* is defined as the angle between two successive displacement vectors **D***i* and **D***i*+1,

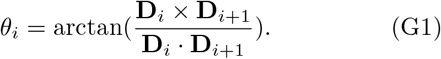

The variance of turning angles is calculated over all turning angles via

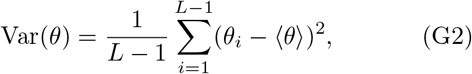

Where 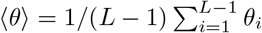.

##### b. End-to-end distance

The absolute value of end-to-end distance is defined as the distance between the first and last monomers of a polymer,

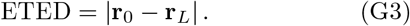

##### c. Radius of gyration

The radius of gyration is defined as the square root of the sum of the diagonal elements of the gyration tensor *S*, which characterizes the spatial distribution of monomers relative to the polymer’s center of mass. The element of the gyration tensor, *Smn*, is defined as

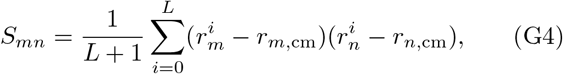

where 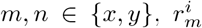 denotes the *m*-th component of **r***i*, and *rm*,cm is the *m*-th component of the center of mass, given by 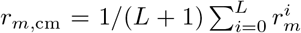. The radius of gyration, *Rg*, is

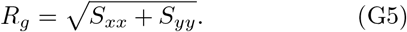

##### d. Mean cosine of turning angles

The mean cosine of turning angles is defined as

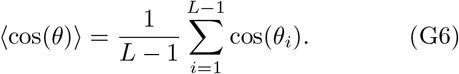

##### e. Mean tangent of turning angles

The mean tangent of turning angles is defined as

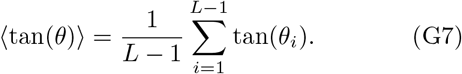

#### 2. Normalized mutual information between important latent features and physical observables

The normalized mutual information values measured between key latent features and relevant physical quantities are presented in Fig. A4. Specifically, we show the NMI values for four of the five most important features (indices 9, 39, 34, 45), excluding the most important one (index 36).

**FIG. A4.**
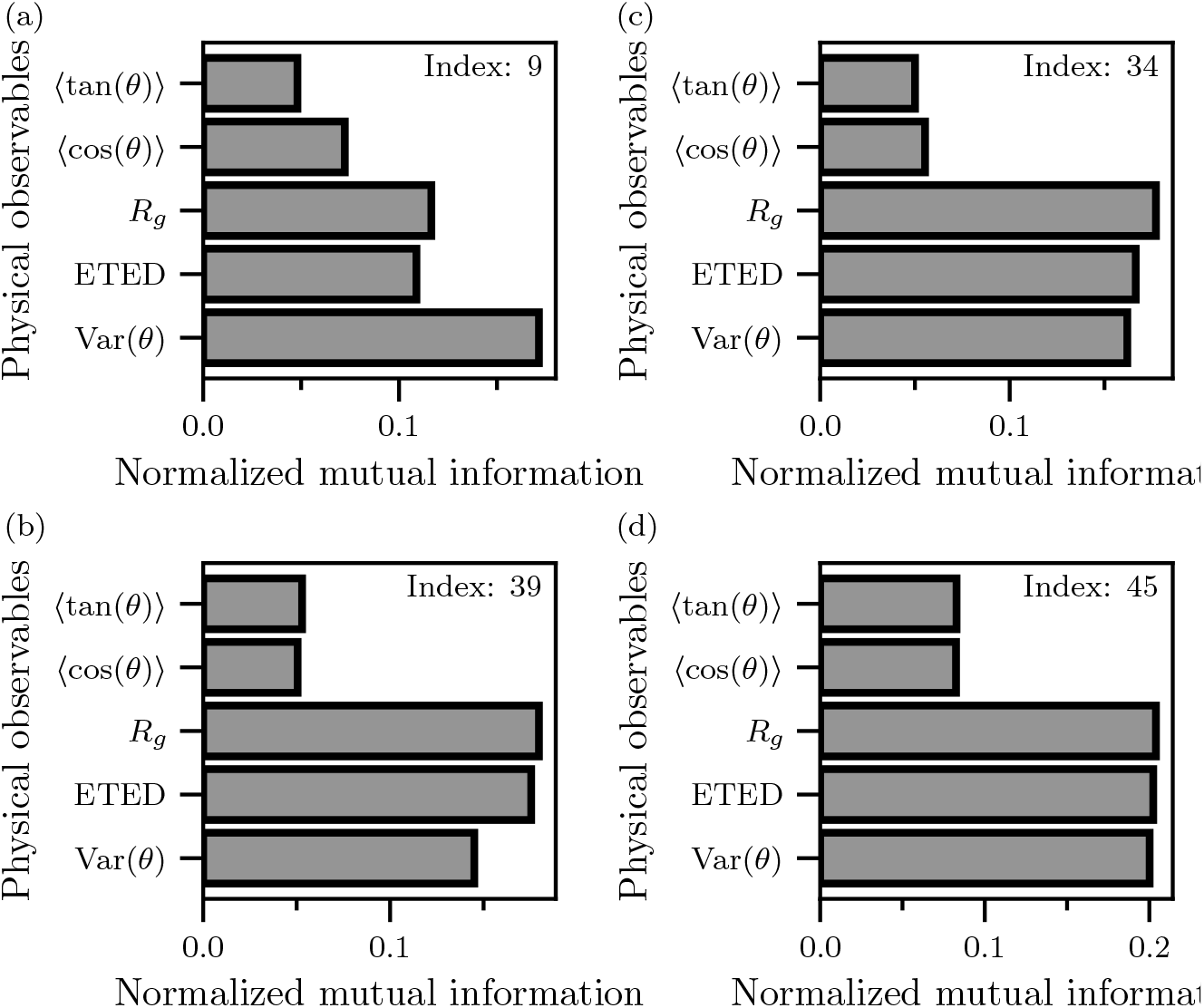
(a-d) Normalized mutual information between important latent features and relevant physical quantities described in Appendices G 1 and G 2.

#### 3. Averaged self-attention matrix

##### a. Definition of averaged self-attention matrix

In the self-attention mechanism, the input sequence (of length *Lw* + 1) to each multi-head attention layer, denoted by *X*IN is linearly projected into three distinct representations: Queries (*Q*), Keys (*K*), and Values (*V*). These projections are computed via

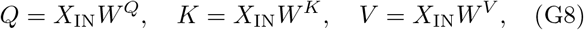

where *W Q, W K*, and *W V* are learnable weight matrices used to generate the query, key, and value representations. Each of these representations is designed to capture dependencies between tokens, which refer to elements in the input sequence. The query encodes the current token’s intent to extract relevant features, while the key encodes the attributes of all tokens that may be attended to. The self-attention matrix, denoted by *A*(*Q, K*), is then computed as the scaled dot product between the queries and keys,

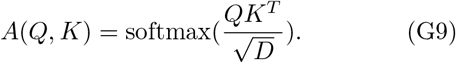

Here, *D* denotes the dimensionality of queries and keys, *T* presents the transpose operation, and the softmax function is applied row-wise across the key dimension. Consequently, the self-attention matrix produced by a single attention head is of size (*Lw* + 1) × (*Lw* + 1). Since each transformer block contains 64 attention heads in our model, a total of 64 self-attention matrices are generated per block. To characterize the average dependency across all heads and blocks of our model, we compute the element-wise average of these matrices, resulting in a single matrix referred to as the averaged self-attention matrix, denoted by *Ā*.

##### b. Visualization of averaged self-attention matrices

The averaged self-attention matrices, *Ā*, are visualized as heatmaps in Fig. A5. Each heatmap is obtained by averaging the averaged self-attention matrices over 1000 polymer samples for a given persistence length.

**FIG. A5.**
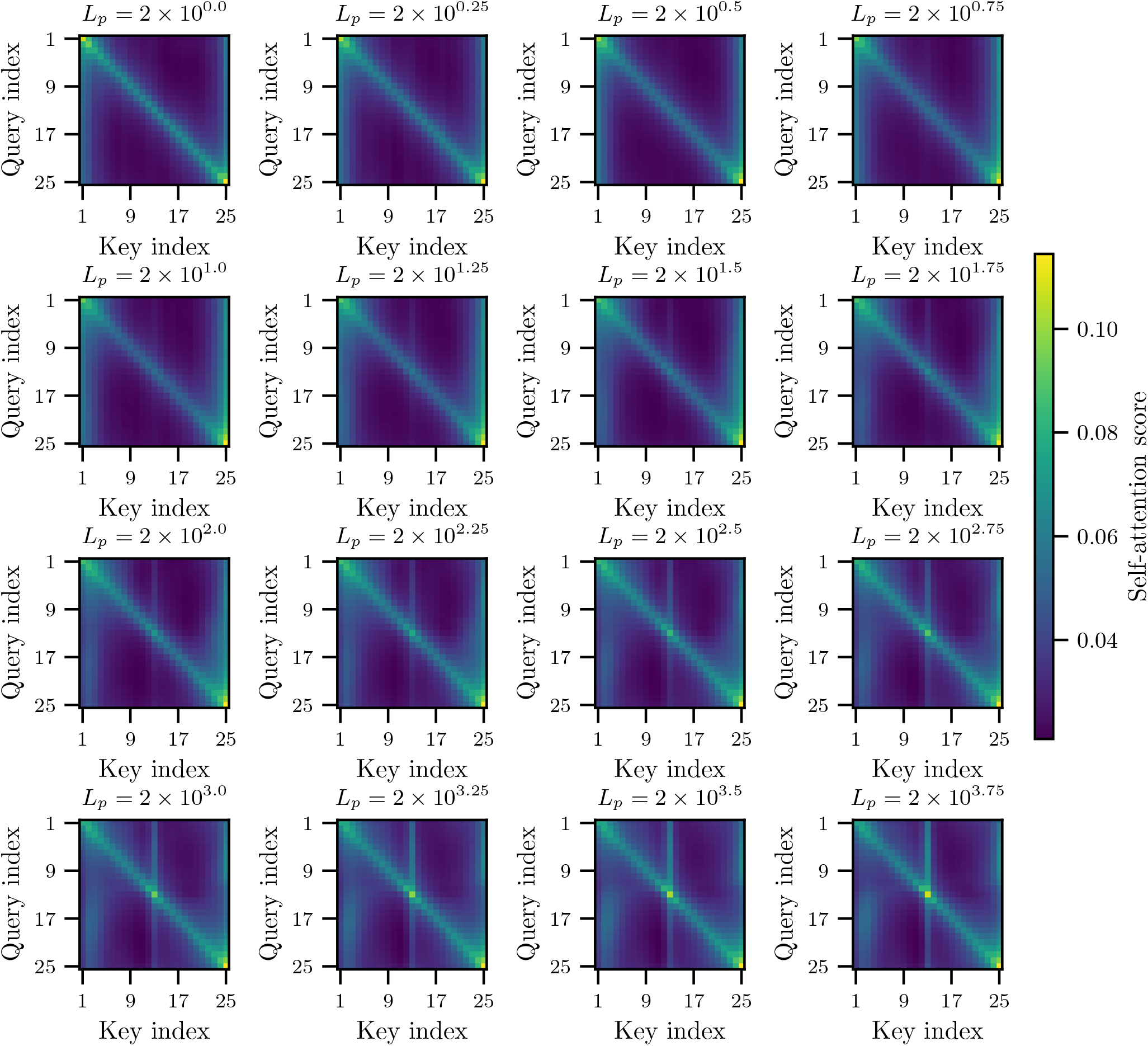
Averaged self-attention matrices for each persistence length, visualized as heatmaps colored by self-attention scores.

## References

[1] J. L. Ross, M. Y. Ali, and D. M. Warshaw, Current Opinion in Cell Biology 20, 41 (2008).

[2] S. Etienne-Manneville, Traffic 5, 470 (2004).

[3] M. Dogterom and G. H. Koenderink, Nature Reviews Molecular Cell Biology 20, 38 (2019).

[4] J. Yan and J. F. Marko, Phys. Rev. Lett. 93, 108108 (2004).

[5] J.-H. Jeon, P. J. Park, and W. Sung, The Journal of Chemical Physics 125, 164901 (2006).

[6] J.-H. Jeon, J. Adamcik, G. Dietler, and R. Metzler, Physical Review Letters 105, 208101 (2010).

[7] D. M. Bangalore, H. S. Heil, C. F. Mehringer, L. Hirsch, K. Hemmen, K. G. Heinze, and I. Tessmer, Scientific Reports 10, 15484 (2020).

[8] H. Kang, M. J. Bradley, W. Cao, K. Zhou, E. E. Grintsevich, A. Michelot, C. V. Sindelar, M. Hochstrasser, and E. M. De La Cruz, Proceedings of the National Academy of Sciences of the United States of America 111, 17821 (2014).

[9] E. M. De La Cruz, J.-L. L. Martiel, and L. Blanchoin, Biophysical Journal 108, 2270 (2015).

[10] A. M. Lorenzo, E. M. De La Cruz, and E. F. Koslover, Soft Matter 16, 2017 (2020).

[11] T. Hawkins, M. Mirigian, M. Selcuk Yasar, and J. L. Ross, Journal of Biomechanics 43, 23 (2010).

[12] A. J. S., R. Padinhateeri, and D. Das, Soft Matter 16, 3125 (2020).

[13] F. Gittes, B. Mickey, J. Nettleton, and J. Howard, Journal of Cell Biology 120, 923 (1993).

[14] C. Bustamante, J. F. Marko, E. D. Siggia, and S. Smith, Science 265, 1599 (1994).

[15] M. Kurachi, M. Hoshi, and H. Tashiro, Cell Motility 30, 221 (1995).

[16] C. Yuan, H. Chen, X. W. Lou, and L. A. Archer, Physical Review Letters 100, 1 (2008).

[17] L. H. Wong, N. A. Kurniawan, H.-P. Too, and R. Rajagopalan, Biomechanics and Modeling in Mechanobiology 13, 839 (2014).

[18] D. Valdman, B. J. Lopez, M. T. Valentine, and P. J. Atzberger, Soft Matter 9, 772 (2013).

[19] A. W. Senior, R. Evans, J. Jumper, J. Kirkpatrick, L. Sifre, T. Green, C. Qin, A. Žídek, A. W. Nelson, A. Bridgland, et al., Nature 577, 706 (2020).

[20] J. Jumper, R. Evans, A. Pritzel, T. Green, M. Figurnov, O. Ronneberger, K. Tunyasuvunakool, R. Bates, A. Žídek, A. Potapenko, et al., Nature 596, 583 (2021).

[21] J. Abramson, J. Adler, J. Dunger, R. Evans, T. Green, A. Pritzel, O. Ronneberger, L. Willmore, A. J. Ballard, J. Bambrick, et al., Nature 630, 493 (2024).

[22] G. Muñoz-Gil, G. Volpe, M. A. Garcia-March, E. Aghion, A. Argun, C. B. Hong, T. Bland, S. Bo, J. A. Conejero, N. Firbas, et al., Nature Communications 12, 6253 (2021).

[23] O. Vandans, K. Yang, Z. Wu, and L. Dai, Phys. Rev. E 101, 022502 (2020).

[24] D. Bhattacharya and T. K. Patra, Macromolecules 54, 3065 (2021).

[25] Q. Gao, T. Dukker, A. M. Schweidtmann, and J. M. Weber, Mol. Syst. Des. Eng. 9, 1130 (2024).

[26] O. Kratky and G. Porod, Journal of Colloid Science 4, 35 (1949).

[27] C. Bouchiat, M. D. Wang, J.-F. Allemand, T. Strick, S. M. Block, and V. Croquette, Biophysical Journal 76, 409 (1999).

[28] W. Sung, Statistical Physics for Biological Matter, Graduate Texts in Physics (Springer, 2018).

[29] K. K. McKayed and J. C. Simpson, Cells 2, 715 (2013).

[30] J. Joshi, S. V. Homburg, and A. Ehrmann, Polymers 14, 1267 (2022).

[31] P. A. Wiggins, T. van der Heijden, F. Moreno-Herrero, A. Spakowitz, R. Phillips, J. Widom, C. Dekker, and P. C. Nelson, Nature Nanotechnology 1, 137 (2006).

[32] H. Ju, H. Skibbe, M. Fukui, S. H. Yoshimura, and H. Naoki, iScience 27, 110907 (2024).

[33] A. Braghetto, S. Kundu, M. Baiesi, and E. Orlandini, Macromolecules 56, 2899 (2023).

[34] Y. Wang, L. Lu, Z. Wang, G. Zhou, and Y. Lu, Macro-molecules 57, 7980 (2024).

[35] A. Vaswani, N. Shazeer, N. Parmar, J. Uszkoreit, L. Jones, A. N. Gomez, L. Kaiser, and I. Polosukhin, Attention Is All You Need (2023).

[36] R. Rombach, A. Blattmann, D. Lorenz, P. Esser, and B. Ommer, High-Resolution Image Synthesis with Latent Diffusion Models (2022), arXiv:2112.10752.

[37] OpenAI, J. Achiam, S. Adler, S. Agarwal, L. Ahmad, I. Akkaya, F. L. Aleman, D. Almeida, J. Altenschmidt, S. Altman, et al., GPT-4 Technical Report (2024).

[38] A. Ott, M. Magnasco, A. Simon, and A. Libchaber, Phys. Rev. E 48, R1642 (1993).

[39] A. Ahsan, J. Rudnick, and R. Bruinsma, Biophysical Journal 74, 132 (1998).

[40] F. Chollet et al., Keras (2015).

[41] Martín Abadi, Ashish Agarwal, Paul Barham, Eugene Brevdo, Zhifeng Chen, Craig Citro, Greg S. Corrado, Andy Davis, Jeffrey Dean, Matthieu Devin, et al., TensorFlow: Large-Scale Machine Learning on Heterogeneous Systems (2015).

[42] I. Tessmer, H. Heil, D. Bangalore, and M. Hashemi, Automated AFM analysis of DNA bending reveals initial lesion sensing strategies of DNA glycosylases (2020).

[43] L. Hauke, A. Primeßnig, B. Eltzner, J. Radwitz, S. H. Huckemann, and F. Rehfeldt, FilamentSensor 2.0: An open-source modular toolbox for 2D/3D cytoskeletal filament tracking (2022).

[44] S. L. Gupton and F. B. Gertler, Science’s STKE 2007, re5 (2007).

[45] M. Chang, O.-c. Lee, G. Bu, J. Oh, N.-O. Yunn, S. H. Ryu, H.-B. Kwon, A. B. Kolomeisky, S.-H. Shim, J. Doh, et al., Science Advances 8, eabj3995 (2022).

[46] P. K. Mattila and P. Lappalainen, Nature Reviews Molecular Cell Biology 9, 446 (2008).

[47] A. Zidovska and E. Sackmann, Biophysical Journal 100, 1428 (2011).

[48] M. M. A. E. Claessens, M. Bathe, E. Frey, and A. R. Bausch, Nature Materials 5, 748 (2006).

[49] H. Takatsuki, E. Bengtsson, and A. Månsson, Biochimica et Biophysica Acta (BBA) - General Subjects 1840, 1933 (2014).

[50] A. Mogilner and B. Rubinstein, Biophysical Journal 89, 782 (2005).

[51] E. Atilgan, D. Wirtz, and S. X. Sun, Biophysical Journal 90, 65 (2006).

[52] N. Leijnse, L. B. Oddershede, and P. M. Bendix, Proceedings of the National Academy of Sciences 112, 136 (2015).

[53] Y. Lan and G. A. Papoian, Biophysical Journal 94, 3839 (2008).

[54] O. Lieleg, M. M. A. E. Claessens, C. Heussinger, E. Frey, and A. R. Bausch, Phys. Rev. Lett. 99, 088102 (2007).

[55] M. Bathe, C. Heussinger, M. M. A. E. Claessens, A. R. Bausch, and E. Frey, Biophysical Journal 94, 2955 (2008).

[56] E. Nogales, Annual Review of Biochemistry 69, 277 (2000).

[57] A. Desai and T. J. Mitchison, Annual Review of Cell and Developmental Biology 13, 83 (1997).

[58] K. M. Liew, P. Xiang, and L. W. Zhang, Composite Structures 123, 98 (2015).

[59] G. J. Brouhard and L. M. Rice, Nature Reviews Molecular Cell Biology 19, 451 (2018).

[60] F. Pampaloni, G. Lattanzi, A. Jońaš, T. Surrey, E. Frey, and E.-L. Florin, Proceedings of the National Academy of Sciences 103, 10248 (2006).

[61] C. P. Brangwynne, F. C. MacKintosh, and D. A. Weitz, Proceedings of the National Academy of Sciences 104, 16128 (2007).

[62] C. Pallavicini, V. Levi, D. E. Wetzler, J. F. Angiolini, L. Benseñor, M. A. Despósito, and L. Bruno, Biophysical Journal 106, 2625 (2014).

[63] B. Mickey and J. Howard, Journal of Cell Biology 130, 909 (1995).

[64] H. Felgner, R. Frank, and M. Schliwa, Journal of Cell Science 109, 509 (1996).

[65] D. Portran, M. Zoccoler, J. Gaillard, V. Stoppin-Mellet, E. Neumann, I. Arnal, J. L. Martiel, and M. Vantard, Molecular Biology of the Cell 24, 1964 (2013).

[66] M. Andreu-Carbó, C. Egoldt, M.-C. Velluz, and C. Aumeier, Nature Communications 15, 2029 (2024).

[67] J. Sohl-Dickstein, E. A. Weiss, N. Maheswaranathan, and S. Ganguli, Deep Unsupervised Learning using Nonequilibrium Thermodynamics (2015), arXiv:1503.03585.

[68] T. Chen, S. Kornblith, M. Norouzi, and G. Hinton, A Simple Framework for Contrastive Learning of Visual Representations (2020), arXiv:2002.05709.

[69] J. Ho, A. Jain, and P. Abbeel, Denoising Diffusion Probabilistic Models (2020), arXiv:2006.11239.

[70] D.-K. Kim, Y. Bae, S. Lee, and H. Jeong, Phys. Rev. Lett. 125, 140604 (2020).

[71] P. Kowalek, H. Loch-Olszewska, L. Laszczuk, J. Opala, and J. Szwabinski, Journal of Physics A: Mathematical and Theoretical 55, 244005 (2022).

[72] B. Zhou, A. Khosla, A. Lapedriza, A. Oliva, and A. Torralba, in 2016 IEEE Conference on Computer Vision and Pattern Recognition (CVPR) (IEEE Computer Society, Los Alamitos, CA, USA, 2016) pp. 2921–2929.

[73] R. R. Selvaraju, M. Cogswell, A. Das, R. Vedantam, D. Parikh, and D. Batra, in Proceedings of the IEEE International Conference on Computer Vision, Vol. 2017-October (Institute of Electrical and Electronics Engineers Inc., 2017) pp. 618–626.

[74] A. Chattopadhay, A. Sarkar, P. Howlader, and V. N. Balasubramanian, in Proceedings - 2018 IEEE Winter Conference on Applications of Computer Vision, WACV 2018, Vol. 2018-January (Institute of Electrical and Electronics Engineers Inc., 2018) pp. 839–847.

[75] T. Chen and C. Guestrin, in Proceedings of the 22nd ACM SIGKDD International Conference on Knowledge Discovery and Data Mining, KDD ‘16 (Association for Computing Machinery, New York, NY, USA, 2016) pp. 785–794.

[76] D. Nagel, G. Diez, and G. Stock, The Journal of Chemical Physics 161, 054108 (2024).

